# Fast and robust identity-by-descent inference with the templated positional Burrows-Wheeler transform

**DOI:** 10.1101/2020.09.14.296939

**Authors:** William A. Freyman, Kimberly F. McManus, Suyash S. Shringarpure, Ethan M. Jewett, Katarzyna Bryc, The 23 and Me Research Team, Adam Auton

## Abstract

Estimating the genomic location and length of identical-by-descent (IBD) segments among individuals is a crucial step in many genetic analyses. However, the exponential growth in the size of biobank and direct-to-consumer (DTC) genetic data sets makes accurate IBD inference a significant computational challenge. Here we present the templated positional Burrows-Wheeler transform (TPBWT) to make fast IBD estimates robust to genotype and phasing errors. Using haplotype data simulated over pedigrees with realistic genotyping and phasing errors we show that the TPBWT outperforms other state-of-the-art IBD inference algorithms in terms of speed and accuracy. For each phase-aware method, we explore the false positive and false negative rates of inferring IBD by segment length and characterize the types of error commonly found. Our results highlight the fragility of most phased IBD inference methods; the accuracy of IBD estimates can be highly sensitive to the quality of haplotype phasing. Additionally we compare the performance of the TPBWT against a widely used phase-free IBD inference approach that is robust to phasing errors. We introduce both in-sample and out-of-sample TPBWT-based IBD inference algorithms and demonstrate their computational efficiency on massive-scale datasets with millions of samples. Furthermore we describe the binary file format for TPBWT-compressed haplotypes that results in fast and efficient out-of-sample IBD computes against very large cohort panels. Finally, we demonstrate the utility of the TPBWT in a brief empirical analysis exploring geographic patterns of haplotype sharing within Mexico. Hierarchical clustering of IBD shared across regions within Mexico reveals geographically structured haplotype sharing and a strong signal of isolation by distance. Our software implementation of the TPBWT is freely available for non-commercial use in the code repository https://github.com/23andMe/phasedibd.

## 1 Introduction

Modern genetic data sets already number in the millions of genomes and are growing exponentially. Inferring the genomic location and length of identical-by-descent (IBD) segments among the related individuals in these data sets is a central step in many genetic analyses. Ideally, IBD estimates can be obtained from phased haplotypes; this means each diploid individual in the data set is represented by two sequences each of which consists of alleles inherited from a single parent. IBD estimates that are phase aware can improve relationship and pedigree inference (Ramstetter et al. 2017, 2018; Williams et al. 2020), enable health and trait inheritance to be traced (Browning and Thompson 2012; Lin et al. 2013; Vacic et al. 2014; Henden et al. 2016; Belbin et al. 2017; Yang et al. 2019; Henden et al. 2019; Finke et al. 2020), and increase the accuracy of many other inferences regarding demographic history and genetic ancestry (Palamara et al. 2012; Ralph and Coop 2013; Palamara and Pe’er 2013; Martin et al. 2018; Pathak et al. 2018; Browning et al. 2018; Naseri et al. 2019c).

Estimating IBD segments is challenging due to not only the size of the genomic data sets but also due to two types of error that break up IBD segments: genotyping and phase switch error (Figure 1). Genotyping error occurs when the observed genotype of an individual is miscalled due to sequencing or microarray errors. Phase switch errors occur when alleles are assigned to the incorrect haplotype within a diploid individual during statistical phasing. Moreover, IBD segments may contain discordant alleles due to mutation or gene conversion since the common ancestor. Together, these errors and discordances may lead IBD inference methods to fragment true long IBD segments into many shorter, erroneous segments on separate haploid chromosomes. Some of these short fragments of IBD may be below the minimum segment length at which IBD inference methods can reliably make estimates. This can then result in an underestimate of the total proportion of the genome that is IBD since short fragments may be erroneously discarded as false IBD. Additionally, the number of IBD segments shared between the two individuals may be overestimated when a fragmented long IBD segment is erroneously identified as several shorter segments.

**Figure 1:**
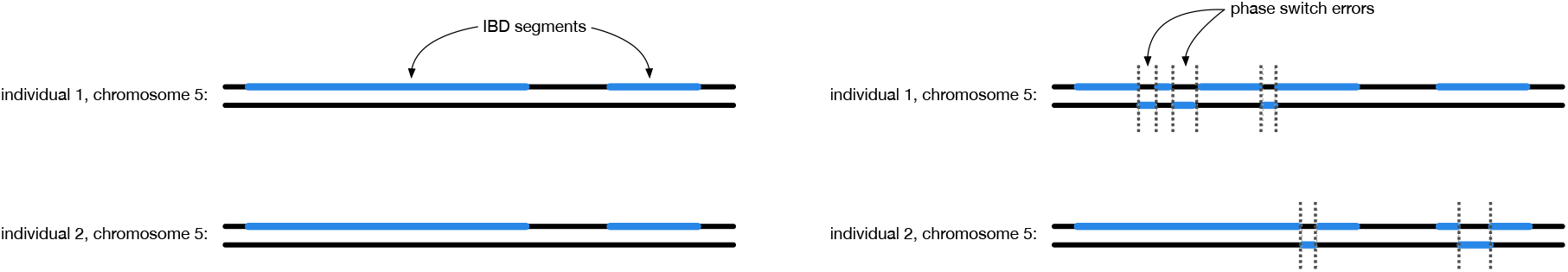
Phase switch errors fragment long IBD segments. *Left:* Two IBD segments (blue) are shared on a single chromosome in two related diploid individuals. *Right:* Phase switch errors (dotted lines) occur at different positions along the chromosome in the two individuals, fragmenting the two true IBD segments into many erroneous short IBD segments. Some of these short fragments of IBD may be below the minimum segment length at which IBD inference methods can reliably make estimates. The discarded fragments can result in an underestimate of the total proportion of the genome that is IBD. Additionally, since each fragment is treated as an individual segment this can result in an overestimate of the number of IBD segments shared between the two individuals.

Here we present the templated positional Burrows-Wheeler transform (TPBWT; see Figure 2), which extends the positional Burrows–Wheeler transform (PBWT; Durbin 2014) to make fast IBD estimates robust to genotype and phasing errors. The TPBWT is an extension of the PBWT with an extra dimension added that masks out potential errors in the haplotypes and extends IBD segments through putative errors. Additionally, we have incorporated within the TPBWT a heuristic that scans patterns of haplotype sharing to identify the location of phase switch errors and correct them. Using haplotype data simulated over pedigrees we explore the speed and accuracy of the TPBWT against other state-of-the-art phase-aware IBD inference approaches in the presence of simulated genotyping and phasing error. For each phase-aware method we compare the false positive and false negative rates of inferring IBD segments of varying lengths. Additionally we compare the performance of the TPBWT against the widely used IBD inference approach described in Henn et al. (2012) that is robust to phasing errors since it uses unphased data. We introduce both in-sample and out-of-sample TPBWT-based IBD inference algorithms and demonstrate their computational efficiency on direct-to-consumer and biobank scale datasets with millions of samples. Finally, we briefly present an empirical analysis that utilizes the TPBWT against the 23andMe database to explore the geographic patterns of haplotype sharing within Mexico. Hierarchical clustering of IBD shared across regions within Mexico reveals geographically stuctured haplotype sharing and a strong signal of isolation by distance.

**Figure 2:**
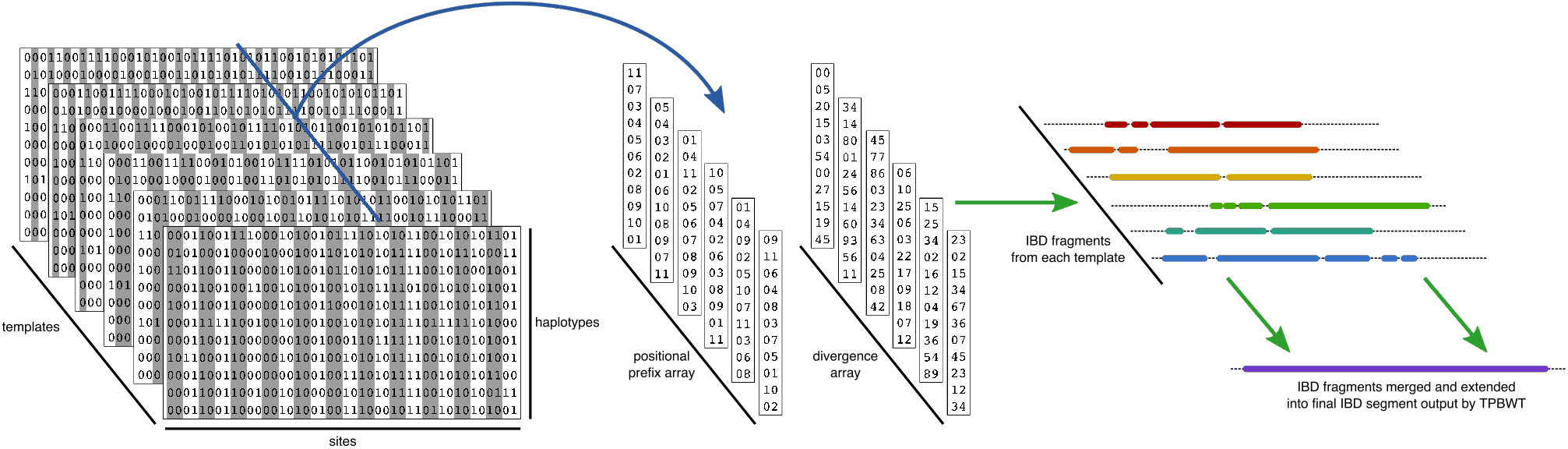
Summary of the TPBWT data structures and IBD inference algorithm. To identify haplotype sharing among a large panel of haplotypes, the TPBWT passes once through a *M* by *N* by *t* three-dimensional structure where *M* is the number of haplotypes, *N* is the number of bi-allelic sites, and *t* is the number of templates. Each template is a pattern at which sites are masked out (shaded out in the figure). During the left-to-right pass through this structure, at each site *k*, two arrays are updated (blue arrow). The positional prefix array *ppa* and the divergence array *div* are both two dimensional arrays of size *M* by *t*. At site *k*, each of the *t* columns of *ppa* and *div* are updated for the templates that are not masked out. Each of the *t* columns in *ppa* contains the haplotypes sorted in order of their reversed prefixes. Similarly each of the *t* columns in *div* contains the position at which matches began between haplotypes adjacent to one another in the sorted order of *ppa*. During the left-to-right pass through this structure, short fragments of IBD shared between haplotypes *i* and *j*, broken up by errors, are identified by each of the *t* templates (green arrows). As these fragments are identified they are merged and extended with one another in the current match arrays 𝒫_*s*_ and 𝒫_*e*_. While merging and extending IBD fragments a heuristic is used to scan for and fix putative phase switch errors. See the main text Section 4 for details.

### 1.1 New Approaches

To detect IBD segments we extend the positional Burrows–Wheeler transform (PBWT; Durbin 2014). Given *M* haplotypes with *N* bi-allelic sites, the PBWT algorithm described in Durbin (2014) identifies identical-by-state (IBS) subsequences of the haplotypes in *O*(*NM*) time. A major limitation of PBWT is that it requires exact IBS subsequence matches with no haplotyping errors or missing data. To reduce sensitivity to error and missing data we introduce the *templated PBWT* (TPBWT) that is inspired by the seed templates used by some short read alignment and homology search algorithms (Ma et al. 2002; Li et al. 2008). The TPBWT identifies IBS subsequences of the haplotypes despite missing data, genotyping, and phase switch errors with only a small linear increase in computational complexity compared to the PBWT.

The TPBWT is robust to error while retaining the speed of the PBWT through two main innovations: (1) the TPBWT adds an extra dimension to the data structures within the PBWT that allows errors to be masked out and haplotype matches to be extended through them, and (2) the TPBWT applies a heuristic that scans for patterns of haplotype sharing to correct putative phase switch errors (Figure 2). To handle genotyping errors, the one-dimensional arrays in the PBWT (described below in the Materials and Methods Section 4) become two-dimensional arrays in the TPBWT. While the PBWT-based algorithm to find IBS sequences passes once through the *N* by *M* two-dimensional haplotype alignment, the TPBWT-based algorithm passes once through a *N* by *M* by *t* three-dimensional structure, where *t* is adjusted to control the method’s sensitivity to error. Each “level” in *t* represents a different template, or pattern, used to mask out sites that may contain errors. During a single pass through this three-dimensional structure, short fragments of IBS, broken up by errors, are identified from each template and then merged and extended. As these fragments of IBS are identified, a heuristic is used to scan for putative phase switch errors by checking the positions of IBS segments on complementary haplotypes. If a phase switch error in one or both of the individuals is found, their phase is corrected and IBS segments previously fragmented by switch errors are merged back together. By identifying and merging IBS fragments while correcting haplotype phasing, the TPBWT achieves better accuracy and computational efficiency than masking out or subsampling sites in multiple independent PBWT runs that are then post-hoc merged. Depending on the degree of sensitivity to error required by the user (determined by parameters described in the text below), the TPBWT has a worst-case time complexity of *O*(*NMt*) or collapses down to the PBWT when *t* = 1. Extensive details on the TPBWT are provided in the Materials and Methods Section 4. Our software implementation is freely available for non-commercial use in the code repository https://github.com/23andMe/phasedibd.

## 2 Results

### 2.1 Performance of TPBWT Versus Other Phase-Aware Algorithms

We compared the performance of TPBWT to other state-of-the-art IBD inference algorithms that use phased data by estimating IBD haplotype sharing within a dataset consisting of haplotypes simulated over pedigrees in which the true IBD shared among individuals was known perfectly. Our simulations included realistic levels of genotyping miscalls and phase switch errors to test how robust each method was to the error found in real data. TPBWT was compared to hap-IBD (Zhou et al. 2019), iLASH (Shemirani et al. 2019), PBWT (Durbin 2014), RaPID (Naseri et al. 2019b), and Refined IBD (Browning and Browning 2013). See Table 2 for parameter settings of the different methods and Section 4.7.4 for a description of the analyses. All methods were run over the same set of simulated haplotypes; see Section 4.7 for details on how the haplotypes were simulated and phased. For each method we examined the IBD inference accuracy, false positive and false negative IBD detection rates, and computational efficiency.

#### 2.1.1 Inference Accuracy

To motivate a systematic comparison of the IBD inference errors from various phase-aware methods, Figure 3 plots the IBD segments estimated by each method and compares them to the true segments for a single randomly selected pair of simulated individuals. Realistic levels of genotyping and phase switch errors were simulated (see Section 4.7). Figure 3 illustrates the nature of the errors from each method; for example, for the single true IBD segment on chromosome 6 the TPBWT correctly estimated a single long IBD segment while the other methods estimated multiple short fragments of IBD: hap-IBD, Refined IBD, and iLASH each estimated two short fragments, RaPID estimated 4 short fragments, and Durbin’s PBWT estimated 6 short fragments. The short fragments of IBD estimated by hap-IBD, Refined IBD, and iLASH covered only a small portion of the true amount of chromosome 6 that was IBD. Note that many of the methods fragmented the true IBD segment at the same locations along the chromosome; these are the locations of phase switch errors. The TPBWT, on the other hand, successfully “stitched” short fragments of IBD together across phase switch and other haplotyping errors to reconstruct the full length of the true IBD segment. Since Durbin’s PBWT was the only method that does not allow for a minimum segment length threshold in genetic distance, it was the only method that detected segments *<* 3 cM; many of those very short fragments filled in gaps between errors and therefore resulted in relatively decent coverage of the genomic regions that were truly IBD but an extreme over estimate in the number of IBD segments. Additionally, Durbin’s PBWT detected very short IBS segments scattered across the genome that were false positive IBD. Note that while the TPBWT appeared to perform the best in terms of accuracy its performance was still far from perfect. For example, all methods including TPBWT erroneously fragmented the single long true IBD segment on chromosome 9 and to varying degrees underestimated the amount of chromosome 9 that was truly IBD (Figure 3). In this case the TPBWT estimated two short segments rather than a single long segment; the other methods all estimated between 7 and 9 short segments.

**Figure 3:**
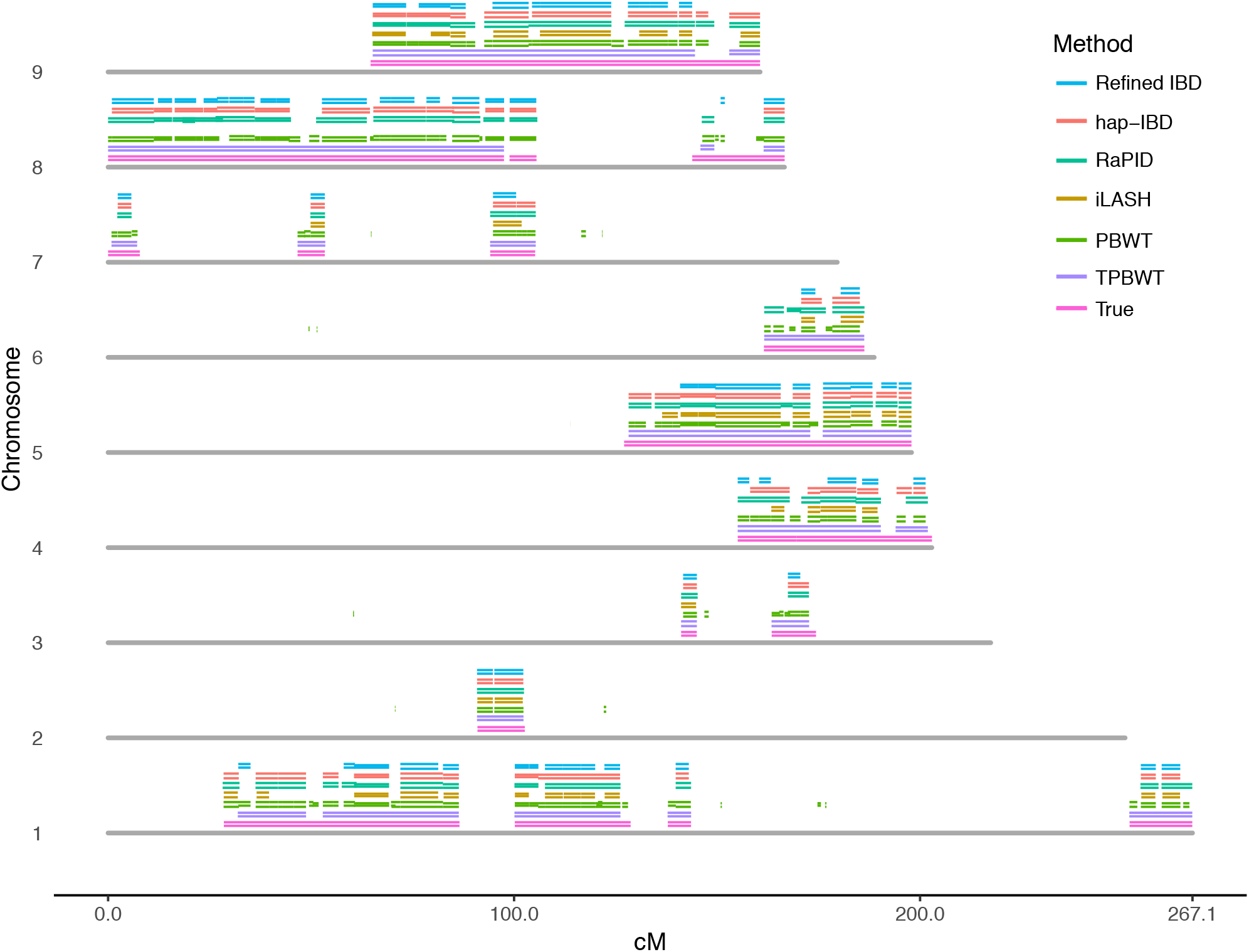
True and estimated IBD segments shared between simulated first cousins. Segments are plotted for chromosomes 1-9 (the other chromosomes were omitted for space considerations). Each chromosome is represented as a grey bar. Above each chromosome are plotted IBD segments; first the true simulated IBD segments (in pink), then segments estimated by each method (in the order depicted in the legend). Each IBD segment is represented by two lines showing their position within each of the two cousins. To a varying degree, phasing errors in either cousin fragmented the IBD segments estimated by each method. For example, all methods including TPBWT erroneously fragmented the single long true IBD segment on chromosome 9. In this case the TPBWT estimated two short segments rather than a single long segment; the other methods all estimated between 7 and 9 short segments. Realistic levels of genotyping and phase switch errors were simulated (see Section 4.7).

To quantitatively compare the performance of the IBD inference methods across a large number of simulations, we focused on their accuracy in estimating two summary statistics: the estimated number of IBD segments shared between two individuals and the estimated proportion of the genome that is IBD between two individuals. These two statistics are particularly informative for downstream analyses such as estimating relatedness and demographic inference. Error in the estimated number of IBD segments shared between simulated relatives is shown in Figure 4. All methods had substantially larger error than TPBWT. The error was highest in closely related pairs that shared long IBD segments; particularly parent–child and siblings. Durbin’s PBWT performed the worst in estimating the number of segments; the true IBD segments were highly fragmented by errors resulting in extreme error, sometimes overestimating by 600 to 800 IBD segments. Error in the estimated percentage of the genome that is IBD in simulated relatives is shown in Figure 5. Here all methods had substantially larger error than TPBWT except Durbin’s PBWT. hap-IBD and Refined IBD had the largest error; on average they underestimated the amount of the genome that was IBD by approximately 10%. The error in all methods was higher in simulated pairs that shared long IBD segments such as parent–child compared to more distant relative pairs such as first cousins. These results confirm the nature of the errors illustrated in Figure 3; compared to the TPBWT, the other methods tested here were highly sensitive to phasing and genotyping errors resulting in estimated IBD segments that were short fragments of the true long IBD segments.

**Figure 4:**
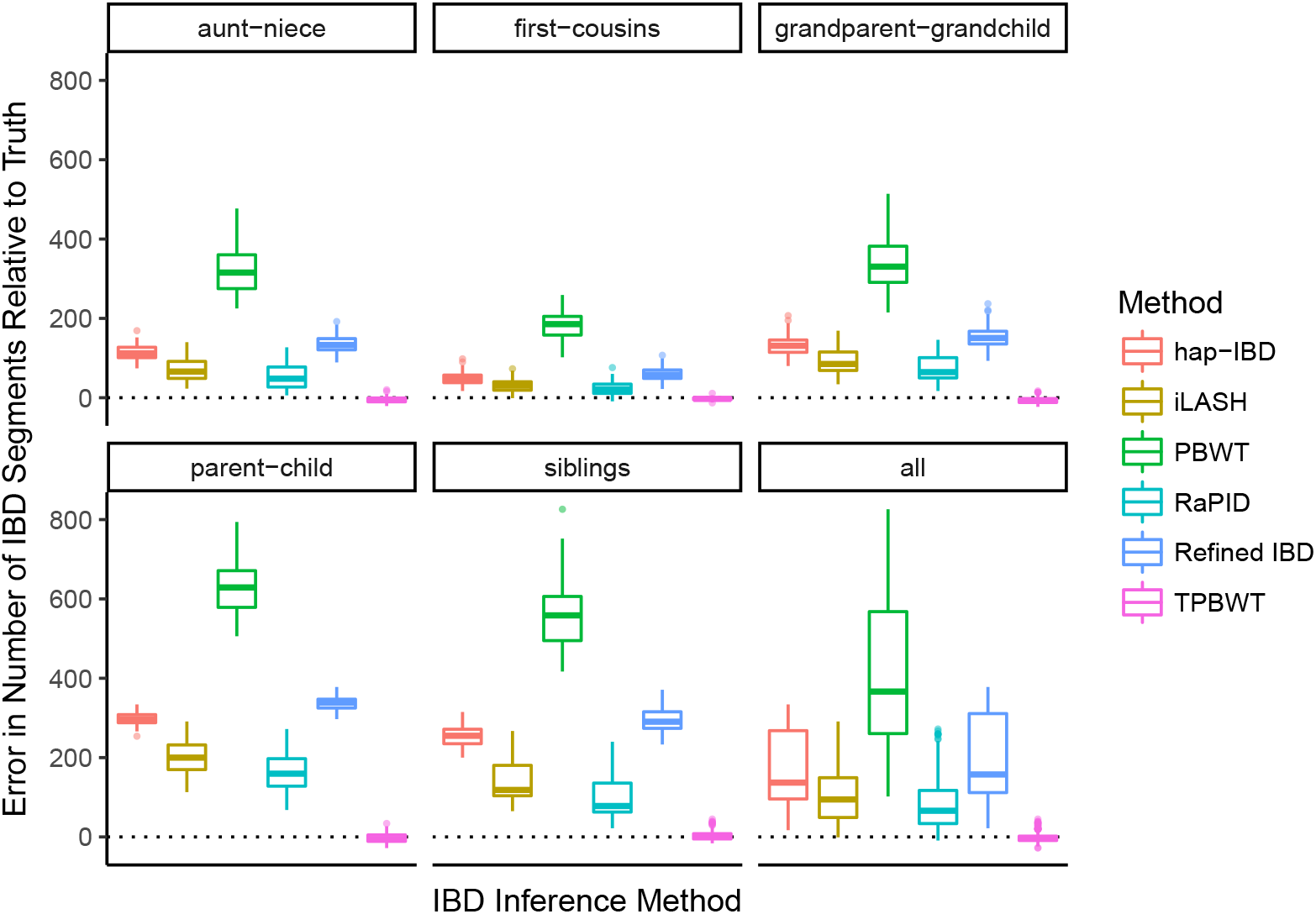
Error in the estimated number of IBD segments shared between simulated relatives. The y-axis represents the number of erroneous IBD segments estimated for a simulated pair of relatives. The error was calculated as (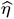 − *η*) where *η* is the true number of IBD segments and 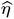 is the estimated number of IBD segments.

**Figure 5:**
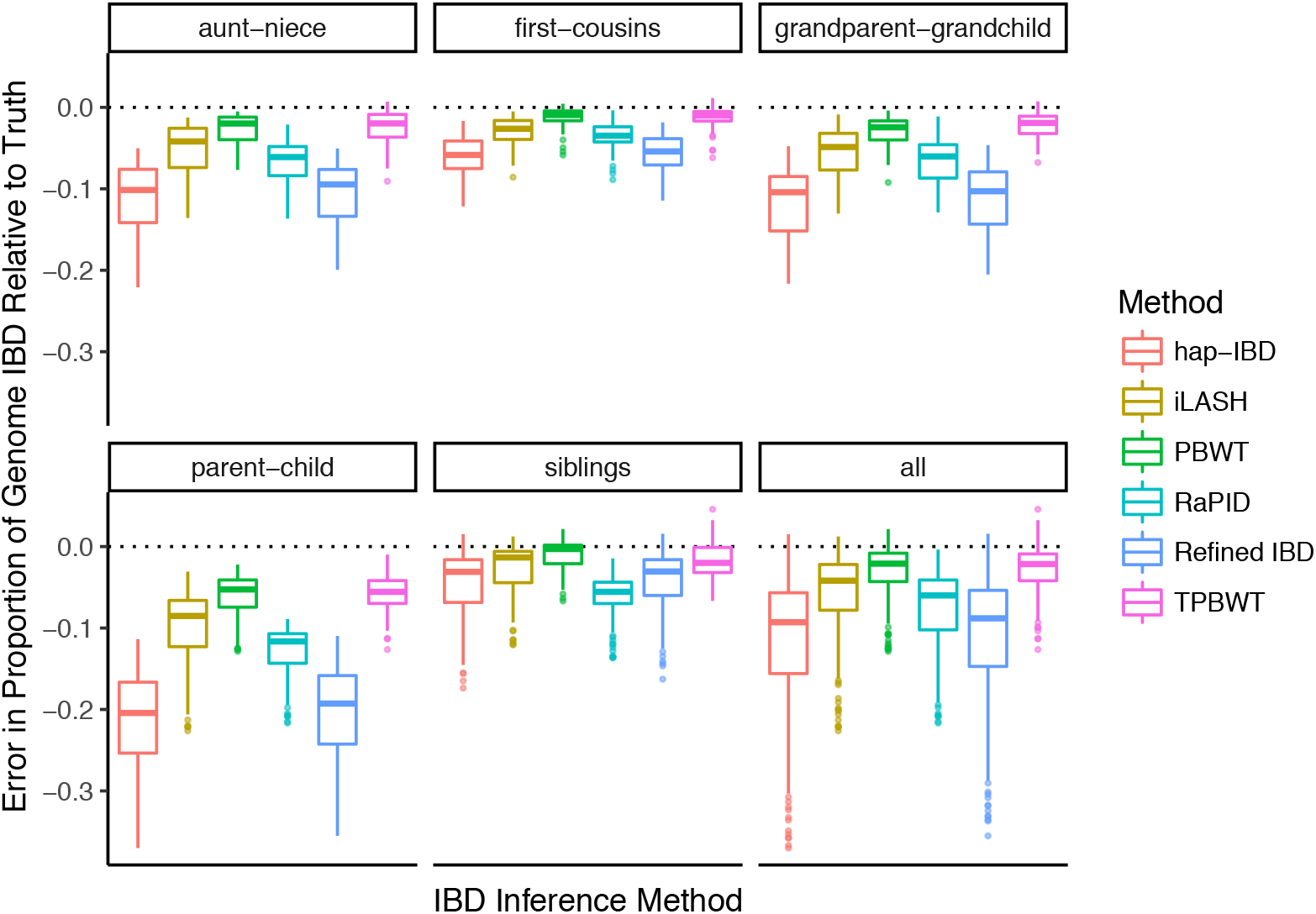
Error in the proportion of the genome estimated to be IBD between simulated relatives. The y-axis represents the proportion of the genome that was erroneously inferred to be IBD for a simulated pair of relatives. The error was calculated as 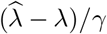 where *λ* is the true total amount of the genome that is IBD, 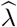 is the estimated amount of the genome that is IBD, and *γ* is the genome length.

#### 2.1.2 False Negative and False Positive Rates

To further characterize the performance of each method we additionally calculated the false positive and false negative rates of inferring IBD. Rates were calculated for bins of IBD segment lengths as described in Section 4.7.4. For IBD segments ≥ 4 cM all methods had very low false positive rates (*<* 0.02; Figure 6). For segments in the smallest bin (3–4 cM) Refined IBD and hap-IBD had the lowest false positive rates (between 0.02 and 0.03). TPBWT, PBWT, and iLASH had false positive rates about 0.04, and RaPID had much higher false positive rates (between 0.4–0.5) compared to all other methods.

**Figure 6:**
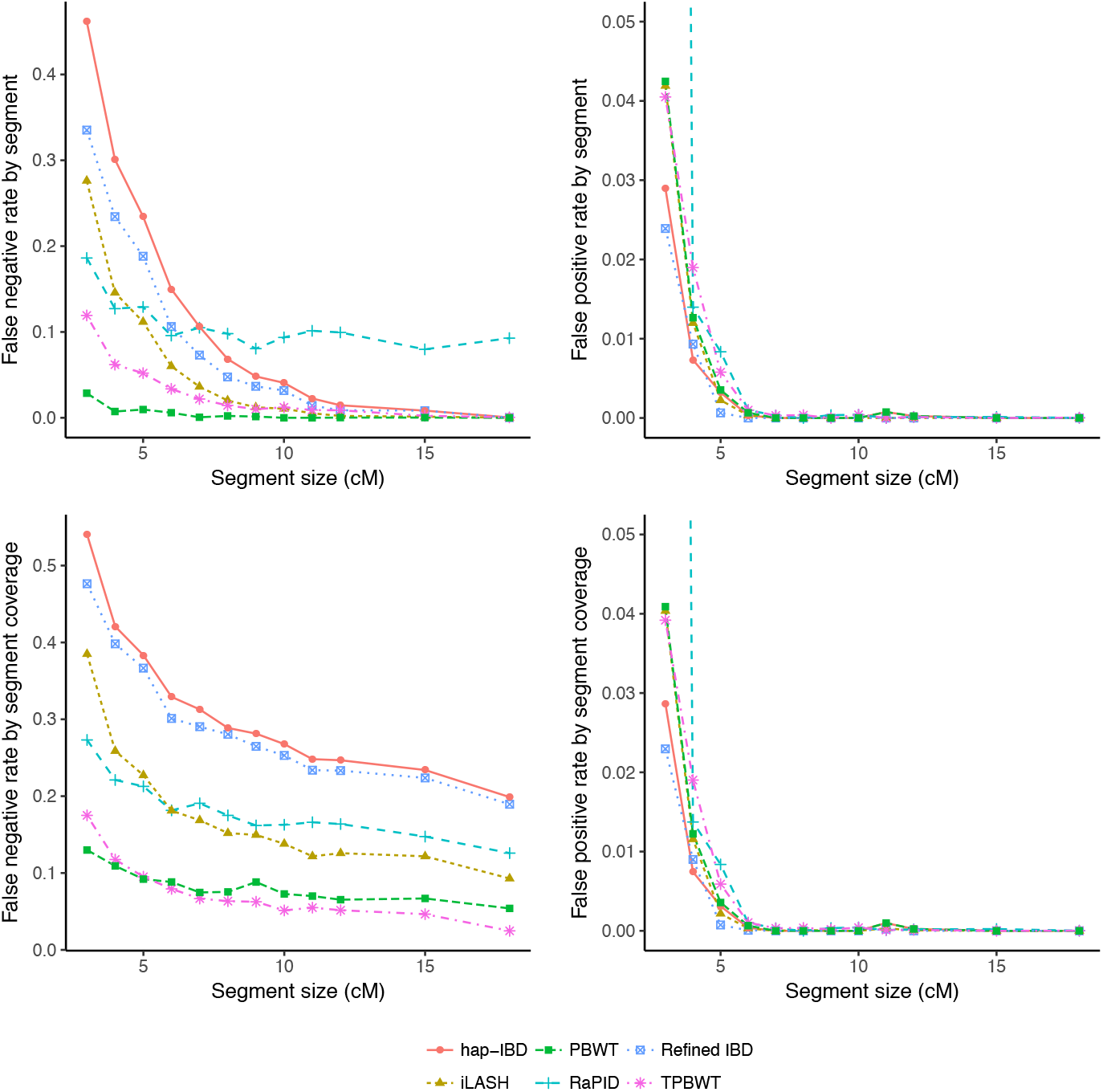
False negative and false positive IBD inference rates. Rates were calculated over simulated data and binned by IBD segment sizes (see main text). The x-coordinate of each point is the lower bound of each size bin (e.g. 3 cM for the 3–4 cM bin). See Table 2 for parameter settings of the different methods. *Left top:* False negative rate by segment is the proportion of true segments in a size bin that do not overlap any estimated segment compared to the total number of true segments in the size bin. *Left bottom:* False negative rate by segment coverage is the proportion of the length of true segments in a size bin not covered by any estimated segment compared to the total length of true segments in the size bin. *Right top:* False positive rate by segment is the proportion of estimated segments in a size bin that do not overlap any true segment compared to the total number of true segments in the size bin. The rate for RaPID in the 3–4 cM bin (cropped out of the plot) was 0.49. *Right bottom:* False positive rate by segment coverage is the proportion of the length of estimated segments in a size bin not covered by any true segment compared to the total length of true segments in the size bin. This rate for RaPID in the 3-4 cM bin (cropped out of the plot) was 0.45.

The false negative rate varied according to how it was calculated (Figure 6). The first false negative rate we compared was calculated as the proportion of true segments in a size bin that did not overlap any estimated segment compared to the total number of true segments in the size bin. Using this rate all methods performed well (approached 0.0) as segment sizes increased except RaPID, which missed approximately 10% of all long segments (*>* 15 cM). However, for short segments the methods varied considerably: in the 3–4 cM range hap-IBD missed over 40% of true IBD segments whereas Durbin’s PBWT missed less than 5% of the segments. For false negatives, all methods performed worse than the TPBWT except Durbin’s PBWT.

The second false negative rate we compared was calculated as the proportion of the length of true segments in a size bin not covered by any estimated segment compared to the total length of true segments in the size bin. Using this rate the TPBWT outperformed all other methods for segments ≥ 6 cM. For segments *<* 6 cM only Durbin’s PBWT outperformed the TPBWT. While the segments estimated by PBWT and TPBWT failed to cover less than 20% of the true segment lengths in the smallest bin (3–4 cM), the other methods failed to cover much higher percentages; in particular hap-IBD and Refined IBD missed approximately 50% of the true segment lengths. For long segments ≥ 18 cM TPBWT was nearly perfect (missed 0%), whereas hap-IBD and Refined IBD missed approximately 25% of the true segment lengths (Figure 6).

#### 2.1.3 Computational Speed

IBD computation runtimes for different methods are shown in Figure 7. Refined IBD and iLASH were at least an order of magnitude slower than the four PBWT-based methods hap-IBD, RaPID, TPBWT, and Durbin’s original PBWT. The four PBWT-based methods all exhibited linear time complexity, while Refined IBD and iLASH took super-linear time. TPBWT was faster than all other methods except Durbin’s PBWT. While iLASH, hap-IBD, and Refined IBD were written as multithreaded programs to take advantage of machines with small numbers of CPU cores the runtimes compared here were for single-threaded operation using a single CPU core. This was done because any of the methods compared here must be parallelized over hundreds of CPU cores using batching approaches to process datasets with millions of samples in reasonable wall clock time (see Section 2.3 and Table 1).

**Table 1:** Compute times for parallelized IBD analyses with large sample sizes. Times are shown for in-sample IBD computes on 1 million individuals, out-of-sample IBD computes on 10k individuals against 1 million, and out-of-sample IBD computes on 10k individuals against 10 million. The first two rows show the compute times measured when IBD was estimated over 42,927 sites of human chromosome 1. The last three rows show those compute times extrapolated to 23 chromosomes with a total of 600k sites. The last row additionally extrapolates the time for an out-of-sample analysis on 1 million to 10 million individuals. CPU time is the sum of the computation time for all compute cores. Wall clock time is the “real” time that the entire analysis took to run.

| IBD Analysis Type | Chromosomes | Number of Samples | Steps Performed | Memory Required (per core) | CPU Cores | CPU Time (minutes) | Wall Clock Time (minutes) |
| --- | --- | --- | --- | --- | --- | --- | --- |
| In-sample | 1 | 1M | 1) In-sample IBD compute and TPBWT-compression for 20 batches of 50k samples | 80 Gb | 20 | 206.4 | 10.3 |
|  |  |  | 2) Out-of-sample comparisons among all compressed batches | 80 Gb | 190 | 2527.0 | 13.3 |
|  |  |  | **Total** |  |  | **2733.4** | **23.6** |
| Out-of-sample | 1 | 10k against 1M | 1) TPBWT-compression of 10k samples | 3.2 Gb | 1 | 1.1 | 1.1 |
|  |  |  | 2) Compare compressed 10k to each compressed 50k batch | 16.0 Gb | 20 | 114.0 | 5.7 |
|  |  |  | **Total** |  |  | **115.1** | **6.8** |
| In-sample | 1–22, X | 1M | 1) In-sample IBD compute and TPBWT-compression for 20 batches of 50k samples | 80 Gb | 460 | 2889.6 | 10.3 |
|  |  |  | 2) Out-of-sample comparisons among all compressed batches | 80 Gb | 920 | 35378.0 | 38.5 |
|  |  |  | **Total** |  |  | **38267.6** | **48.8** |
| Out-of-sample | 1–22, X | 10k against 1M | 1) TPBWT-compression of 10k samples | 3.2 Gb | 23 | 15.4 | 1.1 |
|  |  |  | 2) Compare compressed 10k to each compressed 50k batch | 16.0 Gb | 460 | 1596.0 | 5.7 |
|  |  |  | **Total** |  |  | **1611.4** | **6.8** |
| Out-of-sample | 1–22, X | 10k against 10M | 1) TPBWT-compression of 10k samples | 3.2 Gb | 23 | 15.4 | 1.1 |
|  |  |  | 2) Compare compressed 10k to each compressed 50k batch | 16.0 Gb | 920 | 15960.0 | 17.3 |
|  |  |  | **Total** |  |  | **15975.4** | **18.4** |

**Table 2:** Algorithm parameter values used for the IBD inference methods during the analysis of simulated data. Additionally the same TPBWT parameter values were used for the empirical analysis of geographic patterns of haplotype sharing within Mexico.

| Software | Parameters |
| --- | --- |
| TPBWT | default templates, L<sub>m</sub>=200 L<sub>f</sub>=3.0 |
| PBWT (64c4ffa; Durbin 2014) | -longWithin 200 |
| hap-IBD v1.0 (Zhou et al. 2019) | default options, except nthreads=1 min-output=3 |
| RaPID v1.7 (Naseri et al. 2019b) | -w 3 -r 10 -s 2 -d 3.0 |
| Refined IBD v16May19 (Browning and Browning 2013) | default options, except nthreads=1 length=3.0 |
| iLASH (b26a8fa; Shemirani et al. 2019) | perm_count 12 shingle_size 20 shingle_overlap 0<br>bucket_count 4 max_thread 1 match_threshold 0.99<br>interest_threshold 0.70 min_length 3.0 auto_slice 1<br>cm_overlap 1.4 |

**Figure 7:**
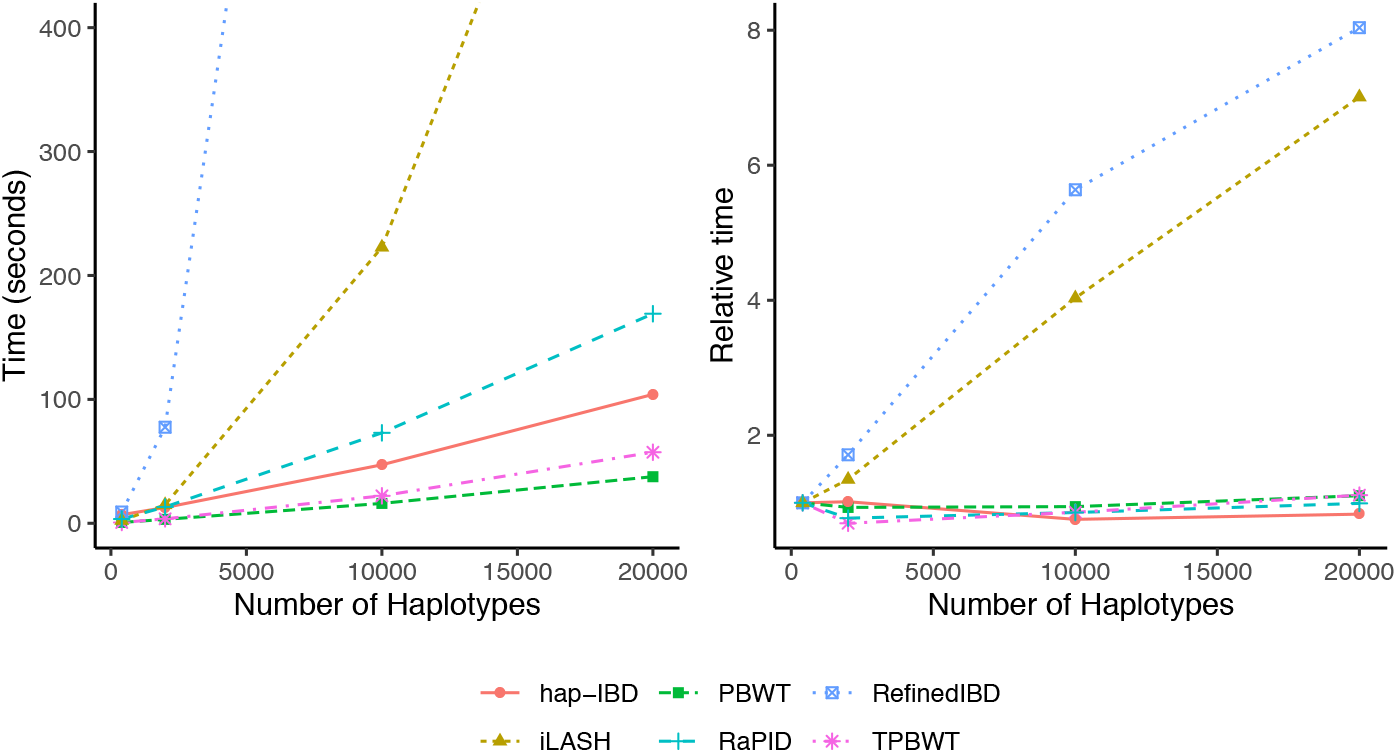
IBD computation runtimes and complexity for different methods. IBD computed for 42,927 SNPs from human chromosome 1. (*Left*) The x-axis is the number of haplotypes analyzed and the y-axis is the time in seconds taken to infer IBD. All methods were run using 1 CPU core. See Table 2 for parameter settings of the different methods. The four PBWT-based methods were at least an order of magnitude faster than the two non-PBWT-based methods; the original Durbin (2014) PBWT was the fastest. Computation times for TPBWT on larger datasets (millions of samples) using the parallel batching approach described in the main text are reported in Section 2.3 and Table 1. (*Right*) Runtime needed to compute IBD for each haplotype in samples sizes of 400 to 20000 haplotypes relative to the time needed to compute IBD for each haplotype in a sample size of 400. Slopes close to zero indicate linear time complexity, positive slopes indicate super-linear time complexity.

### 2.2 Performance of TPBWT Versus a Phase-Free Algorithm

We compared the performance of TPBWT to an IBD inference algorithm that is robust to phasing errors because it uses unphased data. This algorithm was first described in Henn et al. (2012), and was developed independently by Seidman et al. (2020), who called it IBIS. We compared TPBWT to the 23andMe C++ implementation of the IBIS algorithm that was used in Henn et al. (2012), which we refer to here as IBIS-like. The IBIS-like algorithm is known to have a high false positive rate for shorter IBD segments (Henn et al. 2012; Seidman et al. 2020). To account for this while comparing the accuracy of detecting IBD with IBIS-like and the TPBWT, we replicated the trio validation approach used in Henn et al. (2012). For each true IBD segment shared between a child and a distant relative, an overlapping IBD segment between the distant relative and one or more of the child’s parents should also be observed. If this is the case, we labeled the IBD segment “trio validated”. Segments that were not trio validated were either false positive segments in the child or false negative segments in the parents. For bins of IBD segment lengths we calculated the proportion of segments that were trio validated (*h*_mean_) using both TPBWT and IBIS-like, as detailed in Section 4.7.5. Since both Henn et al. (2012) and Seidman et al. (2020) showed that IBIS-like algorithms have high false positive rates for segments *<* 7 cM in length, we used 7 cM has the minimum segment length for the IBIS-like algorithm.

The mean proportion of trio validated segments for bins of IBD segment lengths (*h*_mean_) is shown in Figure 8 panel A. For all bins of IBD segments *>* 6.75 cM TPBWT had trio validation rates of 1.0, which declined to 0.90 for segments in the 3.0–3.25 cM bin. IBIS-like had a trio validation rate of 1.0 for segments in the *>* 14.0 cM bin, which dropped to 0.89 for segments in the 10.0–10.25 cM bin and to 0.42 for segments in the 7.0-7.25 cM bin. This means over half of all the IBD segments in the 7–7.25 cM bin estimated by IBIS-like were either false positive segments in the child or (less likely) false negative segments in the parents. The trio validation rate for TPBWT remained high even for short segments.

**Figure 8:**
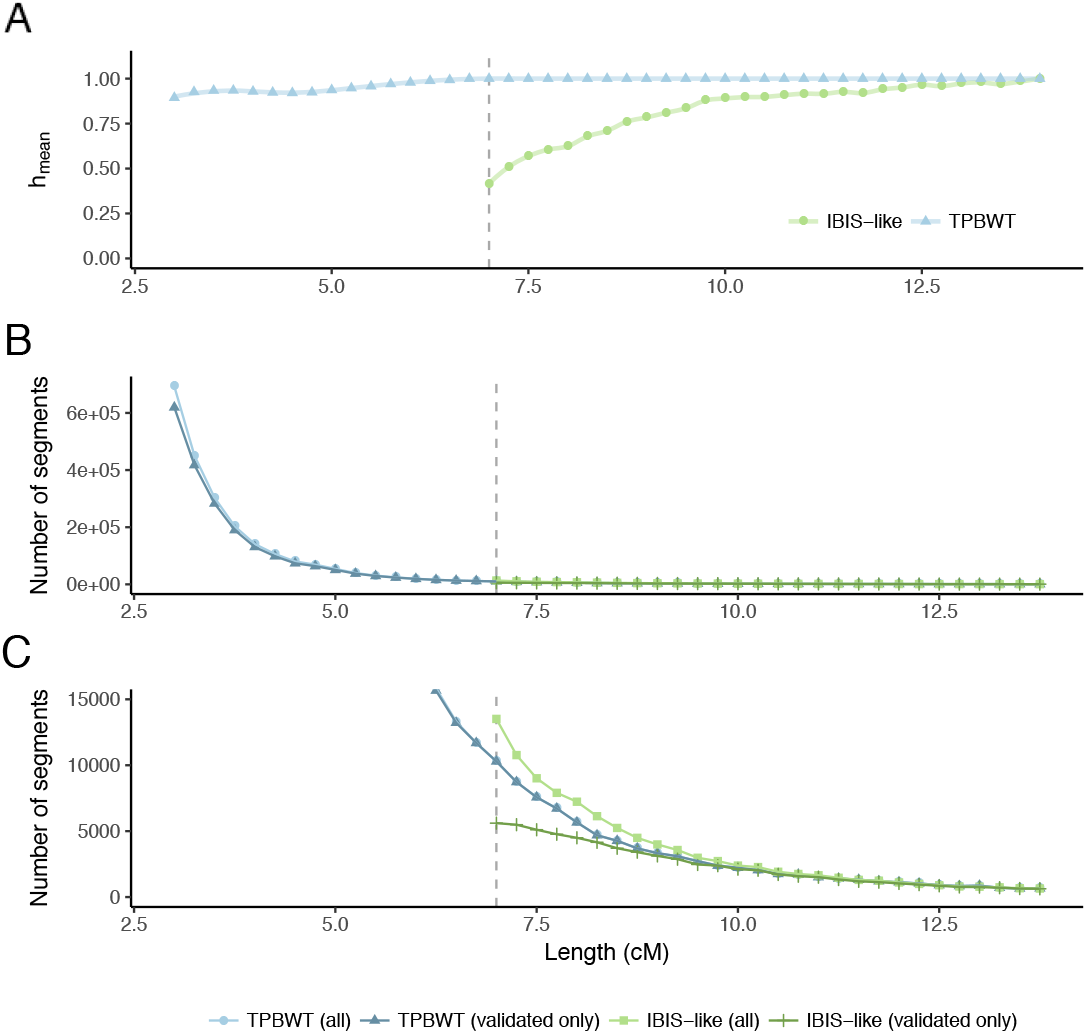
Trio validation test used to compare IBD detection accuracy between TPBWT and the phase-free IBIS-like algorithm. IBD trio validation tests were performed by computing the in-sample IBD among 13,000 individuals (1,000 child-parent trios and 10,000 distant relatives). See main text for details. (*A*) The mean proportion of trio validated segments for bins of IBD segment lengths (*h*_mean_) is shown for TPBWT (blue) and IBIS-like (green). For all bins of IBD segments *>* 6.75 cM TPBWT had trio validation rates of 1.0, which declined to 0.90 for segments in the 3.0–3.25 cM bin. IBIS-like had a trio validation rate of 1.0 for segments in the *>* 14.0 cM bin, which dropped to 0.89 for segments in the 10.0–10.25 cM bin and to 0.42 for segments in the 7.0-7.25 cM bin. (*B*) The number of all segments (trio validated and not trio validated) detected by each method for bins of IBD segment lengths is shown in light green and light blue. The number of trio validated segments detected by each method for bins of IBD segment lengths is shown in dark green and dark blue. The vast majority of the segments detected by TPBWT were *<* 5.0 cM in length, and most of these were trio validated. (*C*) Zoomed in counts of IBD segments reveals that while IBIS-like detected more overall segments 7-8 cM in length than TPBWT, TPBWT detected more trio validated segments 7-8 cM in length than IBIS-like.

The number of segments detected by each method for bins of IBD segment lengths is shown in Figure 8 panels B and C. Using TPBWT a total of 15.5 million segments were detected and using IBIS-like a total of 1.1 million segments were detected. Figure 8 panel B shows that the vast majority of the segments detected by TPBWT were *<* 5.0 cM, and that most of these were trio validated. Using an IBIS-like method this large amount of IBD sharing can not be reliably detected. Figure 8 panel C zooms in on the counts of IBD segments and reveals that while IBIS-like detected more overall segments 7-8 cM in length than TPBWT, TPBWT detected a greater number of trio validated segments 7-8 cM in length than IBIS-like.

### 2.3 Parallelized Performance on Large Cohorts

Table 1 shows both wall clock and CPU runtimes for parallelized IBD analyses on large sample sizes. Wall clock time is the “real” time that the entire analysis took to run. CPU time is the sum of the computation time for all compute cores. The wall clock time taken to compute IBD for 1 million randomly sampled research consented 23andMe customers on chromosome 1 was 23.6 minutes when parallelized across 190 CPU cores. Extrapolated to 23 chromosomes the wall clock time required was 48.8 minutes across 920 CPU cores, well within the capabilities of most HPC cluster facilities.

For large sample size cohorts in biobank or DTC genetic databases out-of-sample IBD computation is an important application. For out-of-sample IBD analyses comparing 10k randomly sampled research consented 23andMe customers to 1 million other customers on chromosome 1 the wall clock compute time required was 6.8 minutes across only 20 CPU cores. Extrapolated to 23 chromosomes and 10 million customers the time needed was 18.4 minutes using 920 CPU cores. These times assumed the haplotypes of the databased cohort (the 10 million individuals) had already been stored as TPBWT-compressed haplotypes. The time needed to TPBWT-compress a set of haplotypes is the same as the time needed to compute their in-sample IBD.

### 2.4 Case Study: Haplotype Sharing in Mexico

Haplotype sharing among 9,517 research consented 23andMe customers who self identified as having all 4 grandparents from a single Mexican state revealed fine-scale population structure within Mexico (Figure 9). Each customer was genotyped on either the 23andMe v4 or v5 microarray chip; after quality control (see Section 4.8) the v4 chip had 453,065 SNPs and the v5 chip had 544,042 SNPs. To minimize the effect of close relatives we excluded any pair of individuals that shared more than 20 cM. Using a single CPU core the IBD compute had a runtime of approximately 20 minutes and revealed 26,606,706 IBD segments across all chromosomes.

**Figure 9:**
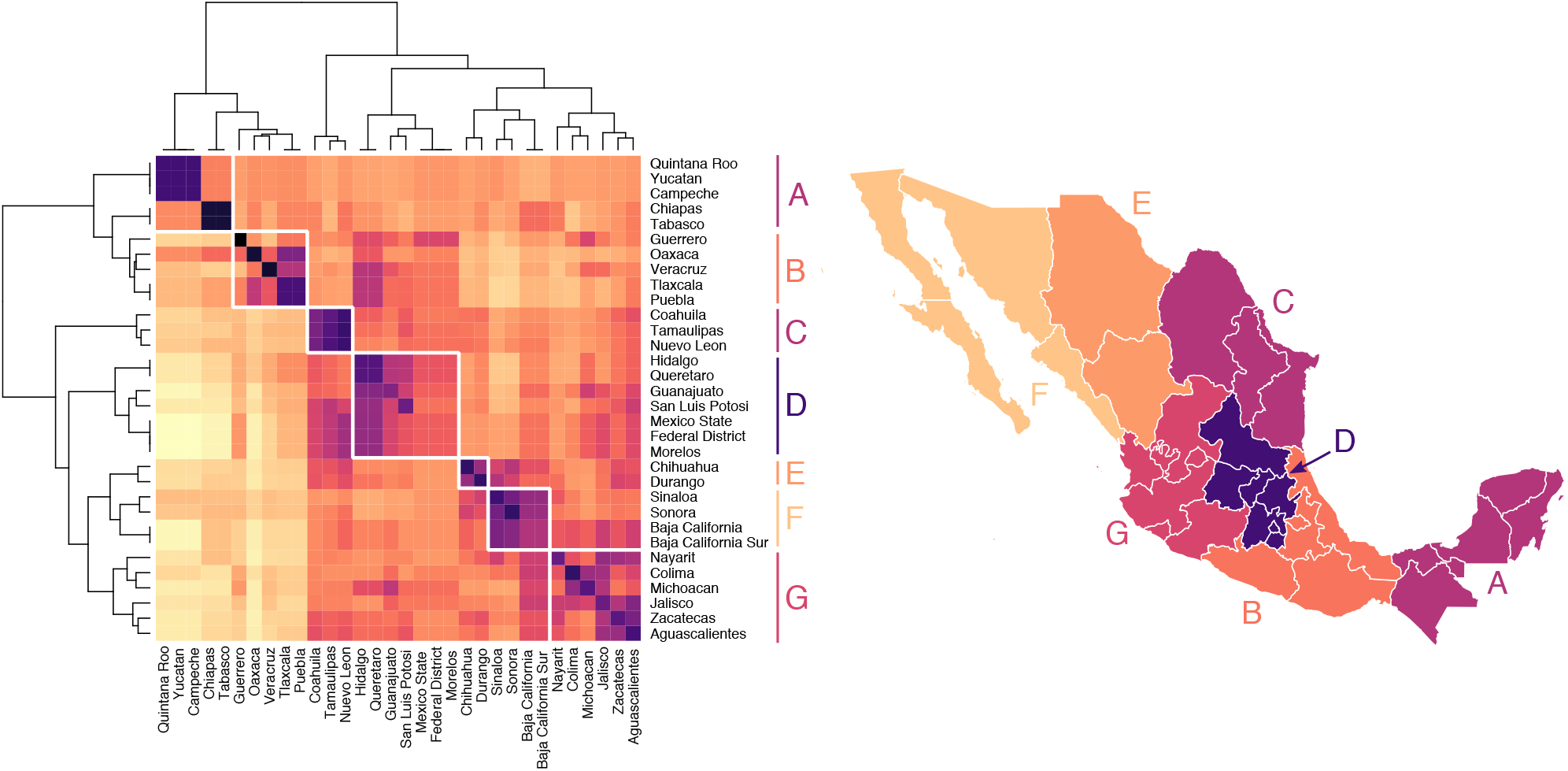
Genetic structure across regions of Mexico. *(Left)* Heatmap showing mean pairwise IBD haplotype sharing across Mexican states. This is the mean sum of the length of IBD segments per pair, where each individual has all 4 grandparents from the same Mexican state. Purple/black represents more sharing, yellow/white represents less sharing. Values are scaled by row. Hierarchical clustering was performed using Ward’s method. The resulting dendrogram shows which states share more IBD, on average, than other states. *(Right)* Population structure is revealed through seven geographic clusters of Mexican states with elevated levels of haplotype sharing (labeled A–G).

Hierarchical clustering of mean pairwise IBD sharing across Mexican states identified geographic clusters of states with elevated levels of haplotype sharing (Figure 9). Our results revealed that IBD sharing among Mexican states decays as geographical distance increases; this is similar to the pattern Martin et al. (2018) found when they clustered the IBD shared among municipal regions of Finland. Our clustering analysis identified clusters of Mexican states that share more IBD on average with one another than with other states; a major limitation of this state-level analysis is that it obscures underlying continuous genetic variation that does not follow state lines. Clustering analysis among Mexican states identified two large clusters of states; one cluster representing the states of the Yucatán peninsula and the southern Mexican states and another cluster representing Mexico City and the central and northern states. Within the southern cluster were two subclusters: a cluster representing the Yucatán peninsula (the states of Yucatán, Quintana Roo, Campeche, Chiapas, and Tabasco) and another cluster representing a group of southern states stretching between the Caribean and Pacific coasts (Guerrero, Oaxaca, Veracruz, Tlaxcala, and Puebla). The northern cluster also consisted of two clear subclusters: a distinct cluster of northeast states (Coahuila, Tamaulipas, and Nuevo León), a cluster of north central states (Chihuahua and Durango), and a cluster of states around the Gulf of California (Sinaloa, Sonora, Baja California, and Baja California Sur). Closely related to the Gulf of California cluster was a cluster of central Pacific coast states (Nayarit, Colima, Michoacán, Jalisco, Zacatecas, and Aguascalientes). The last cluster is in central Mexico surrounding Mexico City (Hidalgo, Querétaro, Guanajuato, San Luis Potosi, México, Federal District, and Morelos).

We found mean pairwise IBD haplotype sharing to be highest within states and among geographically neighboring states (Figure 10). For example, mean IBD shared among individuals with all 4 grandparents from Nuevo León was 13.4 cM, and the mean pairwise IBD shared between individuals with all 4 grandparents from Nuevo León and individuals with all 4 grandparents from neighboring Coahuila and Tamaulipas was 10.9 and 11.9 cM, respectively. In contrast, mean pairwise sharing between individuals with all 4 grandparents from Nuevo León and individuals with all 4 grandparents from Yucatán was 4.8 cM. Similar geographically structured IBD sharing was found throughout Mexico (Figure 10).

**Figure 10:**
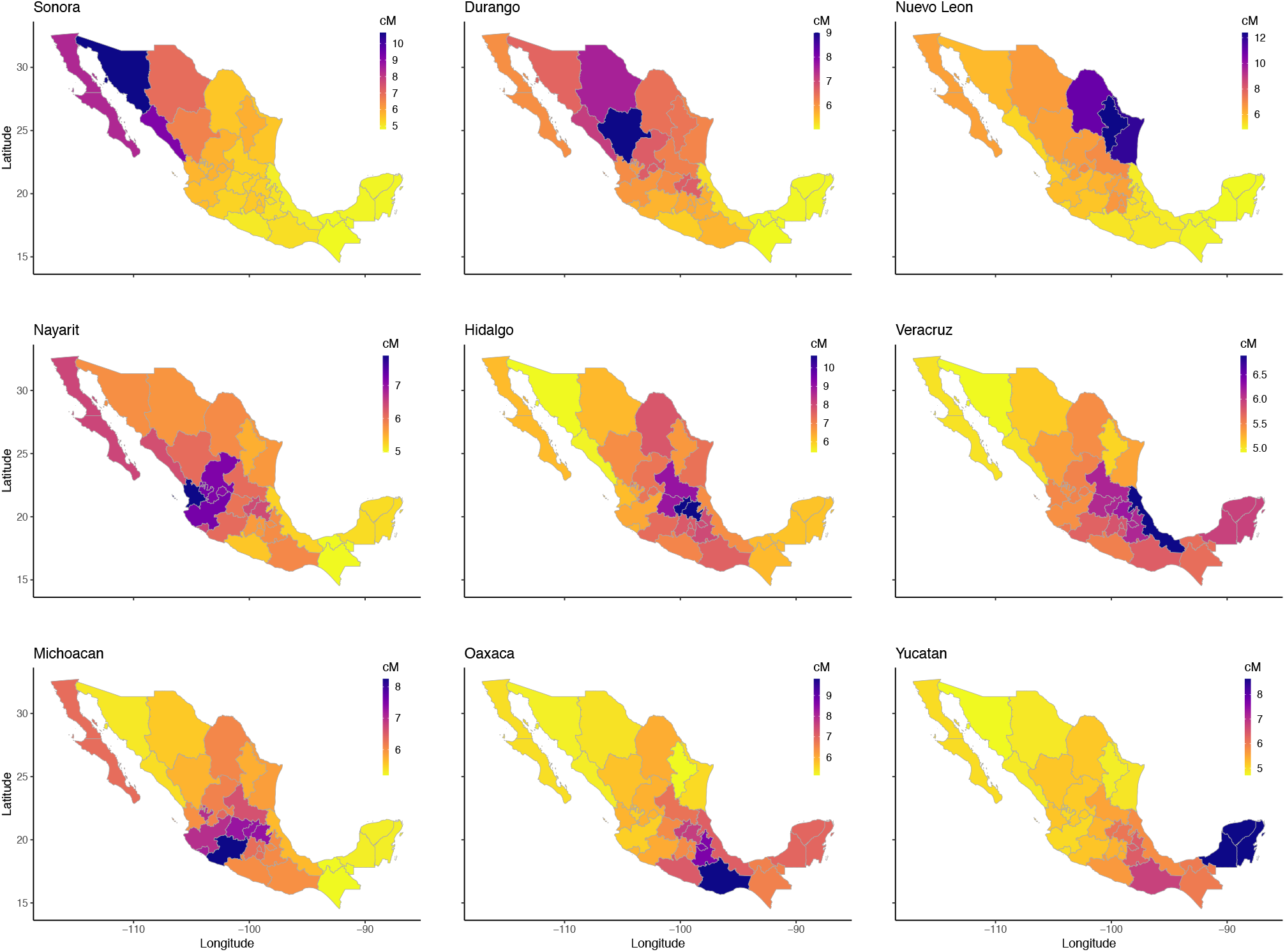
Mean pairwise identical-by-descent (cM) across Mexican states. Within each panel, each Mexican state is colored by the mean pairwise IBD (in cM) shared between individuals with all 4 grandparents from that state and all 4 grandparents from the state specified in that panel. The pairwise IBD is the sum of IBD segments lengths shared between two individuals. Geographically structured IBD sharing was found throughout Mexico. For example, mean IBD shared among individuals with all 4 grandparents from Nuevo León was 13.4 cM. In contrast, mean pairwise sharing between individuals with all 4 grandparents from Nuevo León and individuals with all 4 grandparents from Yucatán was 4.8 cM.

## 3 Discussion

The positional Burrows–Wheeler transform (PBWT; Durbin 2014) was a significant advance in computationally efficient haplotype matching algorithms. Its high sensitivity to error, though, has inspired a number of methods such as RaPID (Naseri et al. 2019b) and hap-IBD (Zhou et al. 2019) that build upon and extend PBWT in an attempt to increase inference accuracy. The templated positional Burrows–Wheeler transform (TPBWT) is similar to these methods in that it extends PBWT to be more robust to haplotyping errors and yet remains highly computationally efficient. However, the TPBWT outperforms these other state-of-the-art phase-aware algorithms due to two primary innovations: (1) the TPBWT adds an extra dimension to the data structures within the PBWT that masks out putative errors and extends haplotype matches through them, and (2) the TPBWT applies a phase correction heuristic that scans for certain patterns of haplotype sharing to identify and correct phase switch errors. The TPBWT’s phase correction method leverages the fact that the patterns of haplotype sharing within large cohorts of samples contain a great deal of information regarding the locations of phase switch errors.

To compare the performance of TPBWT to other state-of-the-art phase-aware IBD inference methods we measured their false negative IBD detection rates in two different ways that together help characterize the nature of IBD inference error. The first, *false negative rate by segment* revealed that most methods did a good job of detecting the presence of a true IBD segment; as long as the true IBD segment was of adequate length then all the methods tested inferred a segment that overlapped the true segment. However, the second *false negative rate by segment coverage* showed that while inferred segments overlapped true segments, often the inferred segments were short fragments that did not adequately cover the entire length of the true segment. In this regard the TPBWT performed substantially better than the other methods compared here. The high false negative rates for the other phase-aware methods were due to estimating highly fragmented IBD segments which led to both an under estimate in the overall percentage of the genome shared as IBD and an over estimate in the number of IBD segments shared. These errors in IBD inference can significantly bias kinship coefficient calculations and negatively impact relationship and pedigree inference.

Our results show that most state-of-the-art phase-aware IBD inference methods performed worse for close relatives compared to more distant relatives; specifically inference accuracy was better for first cousins and aunt-niece pairs compared to parent-child pairs (Figures 4 and 5). Because IBD segments are broken up by meiotic recombination they are expected to be longer for close relatives. Assuming genotyping and phase switch errors are uniformly distributed along the genome, true long IBD segments will on average contain more of these errors than true short IBD segments. This means estimates of long IBD segments are likely to be more negatively impacted by errors compared to estimates of short segments. This can make accurate inference of phase aware IBD among close relatives particularly problematic. Note that Zhou et al. (2019) found much lower false positive rates for hap-IBD than we report in Figure 6. While there are many differences in the datasets used to calculate these rates, one striking difference is that Zhou et al. (2019) evaluated the accuracy of hap-IBD on a dataset consisting of distantly related individuals, whereas our simulation tests focused on closely related individuals. The fact that the negative impact of phase switch errors on the accuracy of phase aware IBD estimates is more severe among closely related individuals may explain hap-IBD’s poorer performance in our tests compared to those by Zhou et al. (2019). We hope that our focus here on accuracy even among closely related individuals will contribute towards methods that make unbiased IBD estimates along the entire spectrum of relatedness.

Another approach to making IBD inference robust to phasing errors is to simply use unphased data. In contrast to the limitations previously discussed with phase-aware IBD methods in accurately identifying IBD among closely related individuals, the major limitation in the accuracy of phase-free IBD methods is in accurately detecting short IBD segments shared among distantly related individuals. 23andMe has used the phase-free algorithm described in Henn et al. (2012) and Seidman et al. (2020) to compute IBD among millions of customers. This approach scans individual’s unphased data for long regions of compatible diplotypes (regions in which two individuals do not have sites with different homozygous genotypes). We show here that using this approach over half of all 7 cM IBD segments estimated were likely false positive segments. If phase-free IBD detection methods such as Henn et al. (2012) and Seidman et al. (2020) are used, downstream quality control filtering of shorter segments should be applied (as is done at 23andMe). Regardless, using these phase-free approaches means that massive amounts of very short (*<* 7 cM) haplotype sharing among distantly related individuals can never be reliably detected. The TPBWT reliably detected segments down to 3-4 cM without any downstream quality control filtering on the segments.

We show here that the TPBWT is not only more accurate and robust to error than other state-of-the-art IBD inference methods but also that it successfully scales to biobank and DTC genetic data sets with millions of samples. One of the most expensive computes for DTC genetic testing companies is calculating the IBD shared between new customers and the entire database of all customers. We presented an example computing IBD for 10,000 new individuals against an existing panel of 10 million individuals. For this compute the TPBWT takes 266.2 CPU hours which, when parallelized appropriately, takes 18.4 minutes of wall clock time.

Additionally, we show that estimates of IBD sharing made using the TPBWT over the 23andMe database can uncover highly granular population structure. Previous studies of population structure in Mexico relied on relatively small sample sizes; data from 66 Mexican-American individuals (Gravel et al. 2013) or 1,000 Mexican individuals (Moreno-Estrada et al. 2014). The scale of the 23andMe database provided a high resolution snapshot of the rich haplotype diversity within Mexico; the IBD sharing among 9,517 research consented 23andMe customers who self identified as having all 4 grandparents from a single Mexican state revealed geographically structured population structure in Mexico. Similar to patterns of IBD sharing at the sub-country level within Finland, our analysis shows that haplotype sharing within Mexico decays with increasing geographic distance (Martin et al. 2018). Expanding upon the IBD estimates presented here with in depth genetic ancestry analyses, as done in Gravel et al. (2013) and Moreno-Estrada et al. (2014), would help increase our understanding of the historical population sizes and migration patterns that led to the rich genetic diversity of Mexico.

For very large biobank and DTC genetic data sets storage and retrieval of previously estimated IBD segments is as large of a computational problem as the initial inference of IBD sharing. Naseri et al. (2019a) presented an algorithm that extends PBWT to compute out-of-sample haplotype sharing between a target and a large panel of pre-indexed haplotypes in constant time. While the method is highly memory intensive, it may be that similar approaches combined with the TPBWT error handling methods introduced here could entirely replace the need to ever store IBD estimates.

Our results highlight the fragility of most phased IBD inference methods; the accuracy of IBD estimates can be highly sensitive to the quality of haplotype estimation. Continued progress on better haplotype phasing methods will undoubtedly help the accuracy of IBD estimates. The two problems are fundamentally linked; indeed both IBD inference methods and phasing methods have benefited from the computational advantages of the PBWT data structure (Loh et al. 2016; Delaneau et al. 2019). Methods that extend PBWT (perhaps incorporating TPBWT-like error handling) to jointly infer IBD and haplotype phase over biobank-scale data sets seem particularly promising. The approach used by the TPBWT to handle missing data is effectively an imputation approach; extending it for more robust imputation would be fruitful. Any TPBWT-based algorithms for phasing and/or imputation could be designed to run directly over TPBWT-compressed haplotypes making large scale reference-based estimates computationally tractable. One unresolved challenge for any PBWT-based inference algorithm is appropriately propagating uncertainty; approaches that integrate probabilistic approaches with the efficiency of PBWT are an exciting way forward.

## 4 Materials and Methods

Inferring IBD segments is challenging primarily due to two types of error that break up IBD segments into short fragments: genotyping and phase switch errors. These errors are particularly problematic when detecting IBD among individuals that are closely related (e.g. first, second, and third degree relatives) since long IBD segments are more likely to be fragmented by these errors. In this work we describe algorithms to compute phase aware IBD segments that are robust to these errors based on a procedure called the templated positional Burrows–Wheeler transform (TPBWT). This is the positional Burrows–Wheeler transform (PBWT; Durbin 2014) with substantial modifications to robustly handle genotyping errors and missing data. Two primary innovations distinguish the TPBWT from the PBWT that increase IBD inference accuracy while retaining the speed of the PBWT. First, the TPBWT adds an extra dimension to the data structures within the PBWT that “templates” or masks the haplotypes, enabling haplotype matches to be extended through errors. This idea of using templates was borrowed from some short read alignment and homology search algorithms (Ma et al. 2002; Li et al. 2008). Second, the TPBWT applies a heuristic that scans for patterns of haplotype sharing to identify the locations of phase switch errors and correct them. Details of each step are given in the sections below.

### 4.1 Templated Positional Burrows–Wheeler Transform (TPBWT)

We will first describe the intuition motivating the TPBWT and then describe the implemented algorithm in detail. One naive approach for extending PBWT to report matching haplotypes that include some error would be to construct multiple replicates of the PBWT data structure. Each of these PBWTs would be built by masking the haplotype alignment using a different repeating template: for example one PBWT could be built that masks out (skips) all the odd positions in the haplotypes, and a second PBWT could be built that masks out all the even positions in the haplotypes. Each PBWT could then be individually swept through identifying exact subsequence matches following algorithm 3 in Durbin (2014). The matching subsequences from each independent PBWT could be merged using a post-hoc algorithm to produce all matching subsequences within the full (unmasked) haplotype alignment. We could modify how sensitive to error this approach is by changing the arrangement and number of templates/PBWTs; in our trivial example of even/odd templates the two templates guarantees that all matches across any two site window will be found as long as there is no more than one error within the window. This is because given any possible location of a single error in the original haplotype alignment at least one of the two PBWT replicates will have that error masked out and therefore still deliver the match correctly. This arrangement of templates would fail if two errors happened to be adjacent to one another in the haplotype alignment. However, for large datasets, the major bottleneck in terms of computational complexity for this naive approach to “templating” the PBWT is the post-hoc algorithm required to merge segments from the PBWT replicates. For every pair of haplotypes sharing IBD the results from each of the individual PBWTs must be scanned through and merged, which has a worst-case time complexity of 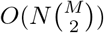. For datasets of non-trivial size (thousands of individuals and greater) much more time will be spent on the post-hoc merging of segments than was spent on the PBWT replicates. Moreover, this naive approach does not share information across the multiple independent PBWT replicates regarding the location of errors. Our goals in developing the TPBWT were to (1) improve accuracy with an algorithm that shares information across “templated” PBWTs so they are no longer independent and, (2) improve the computational efficiency of IBD inference by avoiding the need for post-hoc merging/filtering algorithms.

**Algorithm 1 TPBWT algorithm to find matching subsequences**. The algorithm scans left-to-right along all *N* sites in the haplotype alignment. Here *t* represents the number of templates defined within the templating function *𝒯*and *M* is the number of haplotypes. *𝒯*(*j, k*) denotes the value of *𝒯*for template *j* at position *k*. Additionally *𝒜*_*j,k,i*_ is the allele at position *k* for haplotype *ppa*_*j,k,i*_ and *𝒦* is the next *k* for which *𝒯*(*j, k*) = 1, where *i* is the current template. *m*_0_ and *m*_1_ are temporary lists used to store currently matching haplotypes. *p*_0_, *p*_1_, *d*_0_, *d*_1_ are temporary lists used to assemble *ppa*_*j,k*_ and *div*_*j,k*_. 𝒫_*s*_ and𝒫_*e*_ are each two dimensional arrays that store the current start and end positions of matches between all haplotypes.

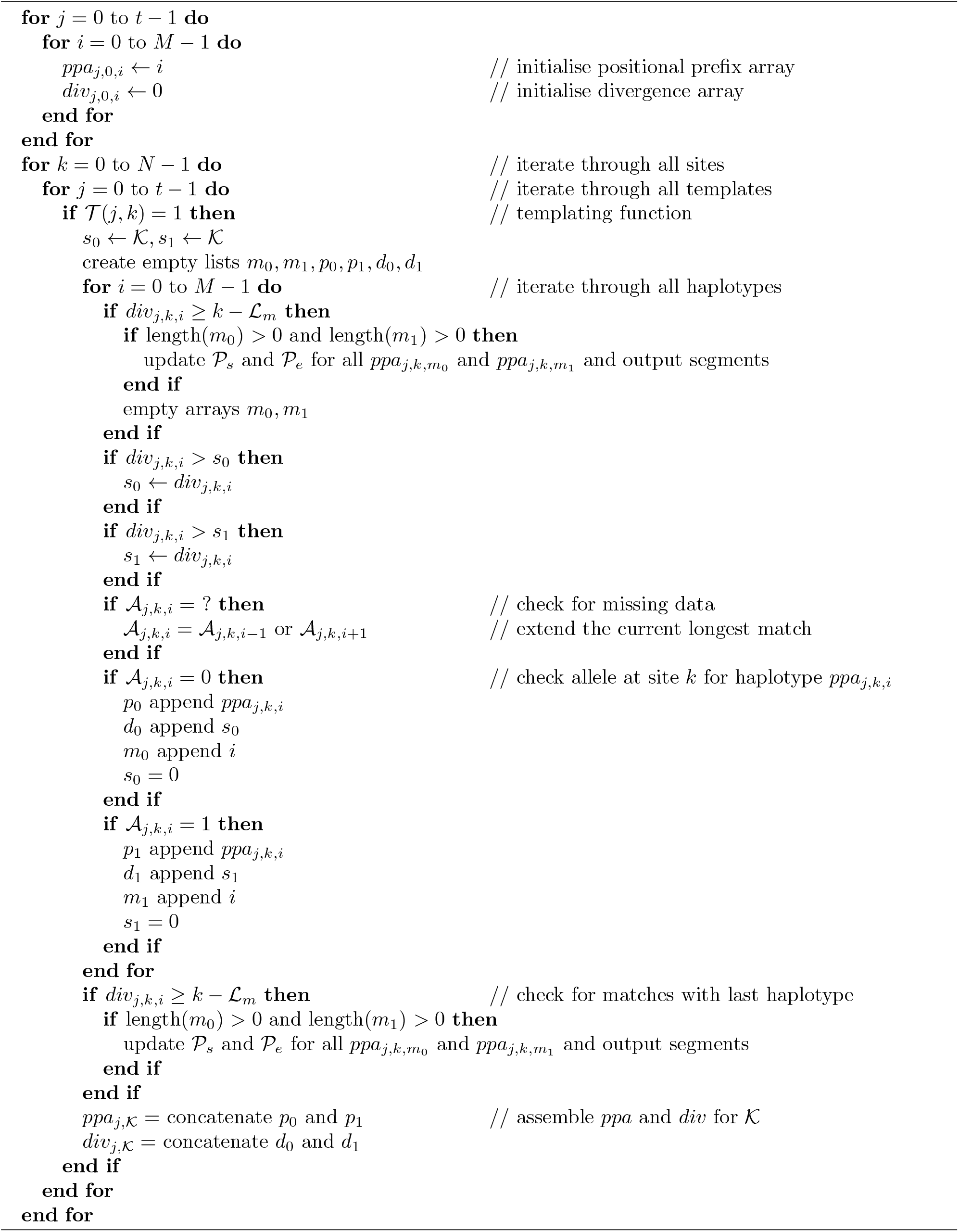

By substantially modifying Durbin (2014)’s PBWT data structure we can utilize the concept of “templating” the PBWT described above to handle errors yet still return subsequence matches in linear time, passing through the data only once and avoiding the need for a post-hoc merging algorithm (see Algorithm 1). In the PBWT, at each position *k* within the haplotype alignment two arrays are constructed: *ppa*_*k*_ the positional prefix array and *div*_*k*_ the divergence array (Durbin 2014). *ppa*_*k*_ is a list of the haplotypes sorted so that their reversed prefixes (from *k −* 1 to 0) are ordered. This ordering ensures that haplotypes that match through position *k −* 1 will end up adjacent to one another in *ppa*_*k*_. The divergence array *div*_*k*_ keeps track of where those matches began, the *i*th element in *div*_*k*_ represents the beginning of the match between the *i*th element in *ppa*_*k*_ and the *i −* 1th element in *ppa*_*k*_. The TPBWT adds an extra dimension to the PBWT that allows errors to be masked out and haplotype matches to be extended through them. The one-dimensional arrays in the PBWT (the positional prefix array and divergence array) become two-dimensional arrays in the TPBWT. While the PBWT-based algorithm to find matching subsequences passes once through the *N* by *M* two-dimensional haplotype alignment, the TPBWT-based algorithm passes once through a *N* by *M* by *t* three-dimensional structure where *t* is the number of templates.

To create the TPBWT and find matching subsequences (Algorithm 1), we construct a separate *ppa*_*j,k*_ and *div*_*j,k*_ for each template *j* used at site *k*. We formalize a set of templates as an indicator function *𝒯*(*j, k*) with the value 0 when the template *j* skips over site *k* and 1 if template *j* processes site *k*. As the haplotype alignment is passed through, *𝒯*(*j, k*) is called for each template *j*; if *𝒯*(*j, k*) is 1 then *ppa*_*j,k*_ and *div*_*j,k*_ are assembled accordingly. If we use a single template and set *𝒯*(*j, k*) to always equal 1, the TPBWT collapses down to the PBWT. When a matching subsequence of at least *ℒ*_*m*_ sites terminates at site *k* under template *j* the start and end positions of the match are stored in auxillary data structures *𝒫*_*s*_ and *𝒫*_*e*_, respectively. *𝒫*_*s*_ and *𝒫*_*e*_ are both *M* by *M* two dimensional arrays in which the position *x, y* holds the start/end positions of the match between haplotype *x* and haplotype *y*. If another subsegment shared between *x* and *y* has already been stored in *𝒫*_*s*_ and *𝒫*_*e*_, we check to see if the new matching subsegment overlaps and possibly extends the existing subsegment. An overlapping subsegment may already have been stored from another template; these two subsegments may be fragments of a single long IBD segment that was broken up by errors. If the two subsegments do not overlap, we check if the old matching segment has a genetic length (in cM) of at least *ℒ*_*f*_ and then report it. The new matching subsegment is then stored in its place. Moreover, we use the arrangement of subsegments within each template to identify possible phase switch errors (described in Section 4.2 below); when a switch error is corrected in one template it immediately affects the output from the other templates. In this way matching subsegments from each template are merged and extended directly through errors with a single pass through the *N* by *M* by *t* three-dimensional structure. Note that *ℒ*_*m*_ is the length of a matching subsegment in the number of sites required to extend a putative IBD segment whereas *ℒ*_*f*_ is the full length in cM for a “good” IBD segment to be called. This formulation allows the user to set *ℒ*_*m*_ to a low value so the algorithm sensitively detects and merges together subsegments of IBD fragmented by error, but only report IBD segments if they extend past a certain genetic length (*ℒ*_*f*_) thus avoiding short runs of IBS to be called false positive IBD.

The TPBWT is more accurate than using multiple independent masked PBWT runs that are post-hoc merged together. Within the TPBWT, each individual “templated” PBWT shares information with the other “templated” PBWTs regarding the location of errors that improves estimates in a way not possible when using multiple independent PBWT runs. As described above, the TPBWT uses arrays *P*_*s*_ and *P*_*e*_ to store fragments of IBS and merge them together, using a heuristic (detailed in Section 4.2) to identify and fix phase switch errors. When a phase switch error is identified using one template, the haplotypes of the individual are swapped in all future sites visited by all templates. Thus, phase switch errors identified in one template effectively modify the ordering of haplotypes in the positional prefix arrays of the other templates; this dependency across templates means that the TPBWT identifies and merges together fragments of IBS that may not have been identified in the first place if using multiple independent PBWTs. Moreover, this means that phase switch errors are fixed consistently throughout the entire cohort; phase switch errors corrected in one individual are consistent across all the IBD that individual shares with all other individuals. This consistency helps ensure that IBD segments can be correctly triangulated within the cohort; if individual A shares a segment with individual B, and individual A shares an overlapping segment with individual C, then individuals B and C should also share an overlapping segment. This is in contrast to phase corrections that are applied pairwise (e.g. Browning and Browning 2011) and so do not guarantee consistency within the cohort.

The TPBWT’s sensitivity to error and speed is modified by the choice of *𝒯*(*j, k*). Depending on *𝒯*(*j, k*), the TPBWT has a worst-case time complexity of *O*(*NMt*) where *t* represent the number of templates defined within *𝒯*(*j, k*); thus the method represents a linear tradeoff between speed and sensitivity to error. In practice genotyping error rates from modern microarrays are low enough that we find an arrangement of six templates is adequate; these templates can be represented as Ø*h*Ø*h, h*Ø*h*Ø, ØØ*hh, hh*ØØ, Ø*hh*Ø, and *h*ØØ*h*, where sites at Ø will be masked out (*𝒯*(*j, k*) = 0) and only sites at *h* will be processed (*𝒯*(*j, k*) = 1). The choice of these six specific templates guarantees that all matches across any given four site window will be found as long as there are no more than two errors within the window. This is because given any possible arrangement of two errors across four sites in the original haplotype alignment at least one of the templates will mask out those errors and therefore still deliver the match correctly. Even with a genotyping error rate as high as 0.001 the probability of three errors within a four site window is 3.996 *×* 10^*−*9^ (assuming error independence). Using this set of templates, the TPBWT has a computation time of about half *O*(*NMt*) because *N* becomes *N/*2 since each template only processes 2 out of every 4 positions in the alignment. More templates could be utilized to ensure matches across longer windows; indeed 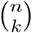 templates are required to ensure all matches across windows of size *n* with no more than *k* errors per window. Similarly with fewer templates the algorithm will run more quickly but be more sensitive to error; when *t* = 1 and *𝒯*(*j, k*) is set to always equal 1 the TPBWT collapses down to the PBWT.

The accuracy of the algorithm with a given set of templates depends on the density of sites and their informativeness in the dataset. For example, consider the case in which a single template *h*ØØØØØØØØØ is utilized to detect IBD; in this case only one tenth of the data is considered when identifying matching subsequences. This choice of templates may provide adequate performance for data with a very high density of informative sites but may negatively affect performance when there is a low density of informative sites. In this case the IBD segments that are correctly identified may be erroneously lengthened and there may be a much higher false positive rate in the IBD segments detected.

Our TPBWT is further detailed as pseudocode in Algorithm 1. The algorithm requires 4 parameters: (1) *𝒯*(*j, k*) which defines the number and arrangement of templates to be used, (2) *ℒ*_*m*_ is the minimum number of sites that a subsegment must span within the haplotype alignment to be merged and extend other subsegments, (3) *ℒ*_*f*_ is the final minimum length (in cM) that a segment must have to be reported by the algorithm, and (4) *M*_*t*_ is the maximum length of a run of missing sites to extend a match through. The algorithm handles missing data by extending the current longest match; implicitly imputing the missing sites using haplotype matching. For template *j* at site *k* the longest matching haplotype to haplotype *ppa*_*j,k,i*_ will be either *ppa*_*j,k,i−*1_ or *ppa*_*j,k,i*+1_, so if missing data in *ppa*_*j,k,i*_ is encountered we simply assume the haplotype continues to extend the longest match. Note that the “imputation” performed here is local by template; *ppa*_*j,k,i−*1_ and *ppa*_*j,k,i*+1_ may differ for each template *j* and so the allele “imputed” may differ for each template. Matches are extended for *M*_*t*_ consecutive missing sites after which they are terminated. Note that the way the algorithm extends matches through runs of missing data *M*_*t*_ sites in length is ommitted in Algorithm 1 for space considerations. One additional detail is not shown in Algorithm 1; after passing through all sites in the haplotype alignment we loop through the haplotypes once last time to report any “trailing” matches (matches that extend all the way through the end of the haplotypes). At this point any matches left in *𝒫*_*s*_ and *𝒫*_*e*_ of length *ℒ*_*f*_ or greater are now reported.

### 4.2 Phase Correction Within the TPBWT

As described above, the TPBWT handles haplotype error (miscalls) and missing data. It is also robust to “blip” phase switch errors in which the phase at a single site is swapped. However, phase switch errors spaced out along the chromosome will cause long regions of the haplotypes to be swapped and fragment IBD segments as illustrated in Figure 1. To handle these errors the TPBWT applies a phase correction heuristic that scans for certain patterns of haplotype sharing to identify and correct phase switch errors (see Figure 11). Note that for haploid data sets such as human male sex chromosomes this heuristic can be turned off. Large cohorts of samples have patterns of haplotype sharing that are highly informative regarding the location of phase switch errors. The phase switch errors in an individual will fragment all IBD segments shared with that individual at the position of the switch error. Each IBD segment that spans the switch error will be broken into two fragments at the position of the error: these fragments will be on complementary haplotypes within the individual with the error and yet remain on the same haplotype within the other individual. For some closely related pairs (parent–child) this pattern of haplotype sharing may be the result of actual recombination patterns, however for the vast majority of more distantly related individuals the pattern can be used to identify phase switch errors.

**Figure 11:**
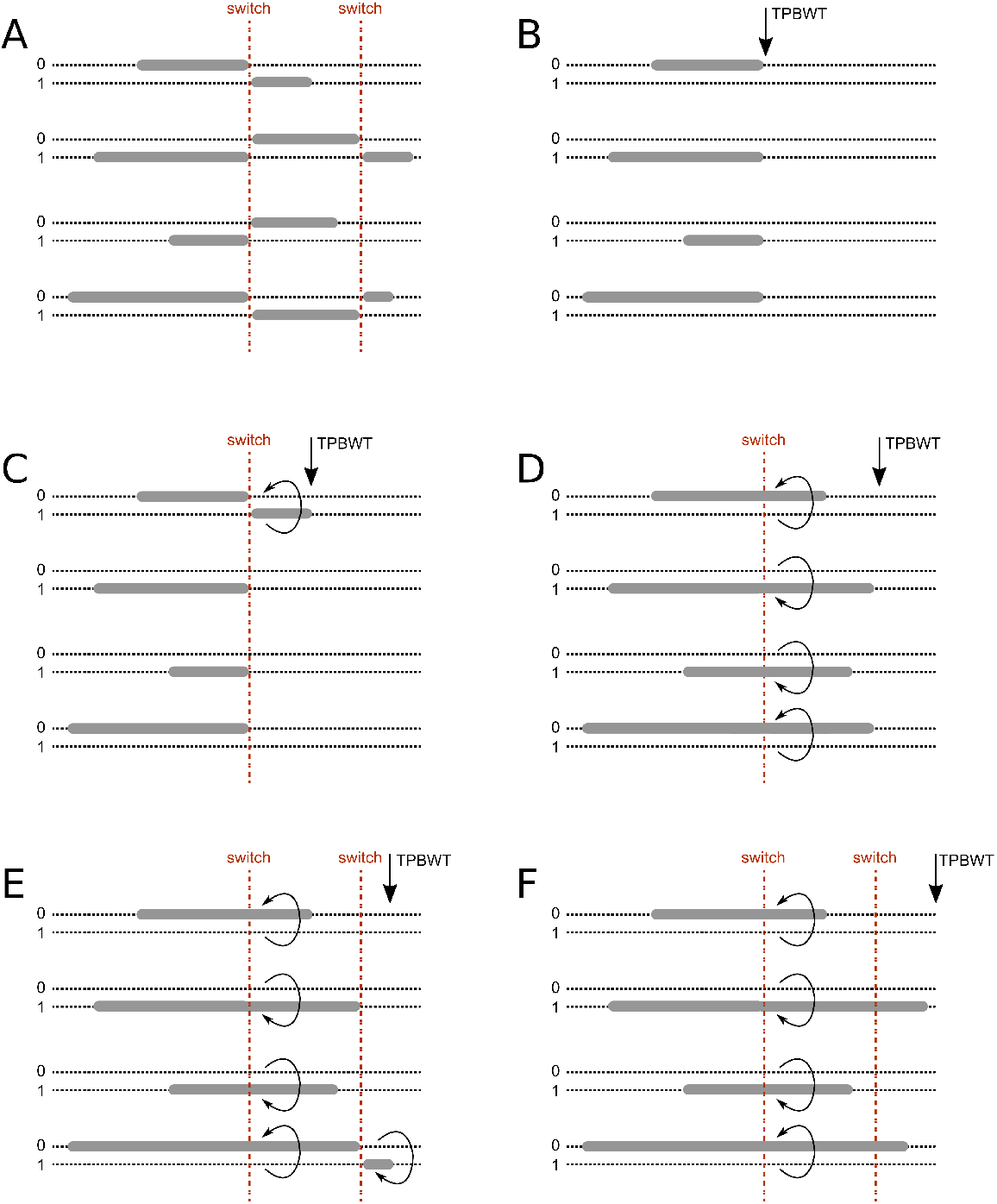
Diagram of the TPBWT phase correction heuristic. As the TPBWT sweeps along the haplotypes identifying IBS matches it uses a heuristic to identify and fix putative phase switch errors. (*A*) The two haplotypes (0 and 1; dotted lines) of a focal person and the IBD segments (grey bars) they share with four other individuals in the haplotype alignment are plotted. The focal person has two phase switch errors (red dashed lines) that break up long IBD segments. (*B*) As the TPBWT scans left to right along the chromosome, it keeps track of IBD segments shared among all pairs of individuals. When a phase switch error is encountered in the focal person all IBD segments shared with the focal person are fragmented at the position of the switch error. (*C*) As the TPBWT continues to scan left to right, another IBD segment is found. If the new segment begins near the end of all the old segments but on the complementary haplotype of the focal person (considering the possible scenarios in Figure 12), then the TPBWT infers a phase switch error to have occurred. (*D*) Since a phase switch error is inferred within the focal person, the focal person’s haplotypes are now swapped so new IBD segments now merge and extend the fragments on the complementary haplotype that were broken up by the phase switch error. (*E*) When the arrangement of IBD segments on the complementary haplotypes again suggests another phase switch error has been encountered the algorithm stops swapping the focal person’s haplotypes. (*F*) The TPBWT continues to the end of haplotypes after successfully identifying phase switch errors and “stitching” IBD fragments back into the correct long IBD segments.

As the TPBWT scans left to right through the haplotype alignment finding new IBD segments it keeps track of previously found IBD segments shared among pairs of haplotypes in *𝒫*_*s*_ and *𝒫*_*e*_. When a new segment shared between two individuals *P* and *Q* is found to be adjacent to an existing segment (either slightly overlapping or with a small gap between them) there are a number of possible scenarios (Figure 12). If the new segment is on the same haplotypes as the existing segment, then possibly the two segments are fragments of a longer segment broken up by error and should be merged. If the new segment begins near the end of the existing segment and the new segment is *not* on the same haplotypes as the old segment, then possibly there was a phase switch error in both individuals. If the new segment begins near the end of the existing segment and the new segment is on the same haplotype as the existing segment in individual *P* but the complementary haplotype in individual *Q*, then possibly there was a phase switch error in individual *Q*. And of course, the opposite pattern could exist suggesting a phase switch error in individual *P*.

**Figure 12:**
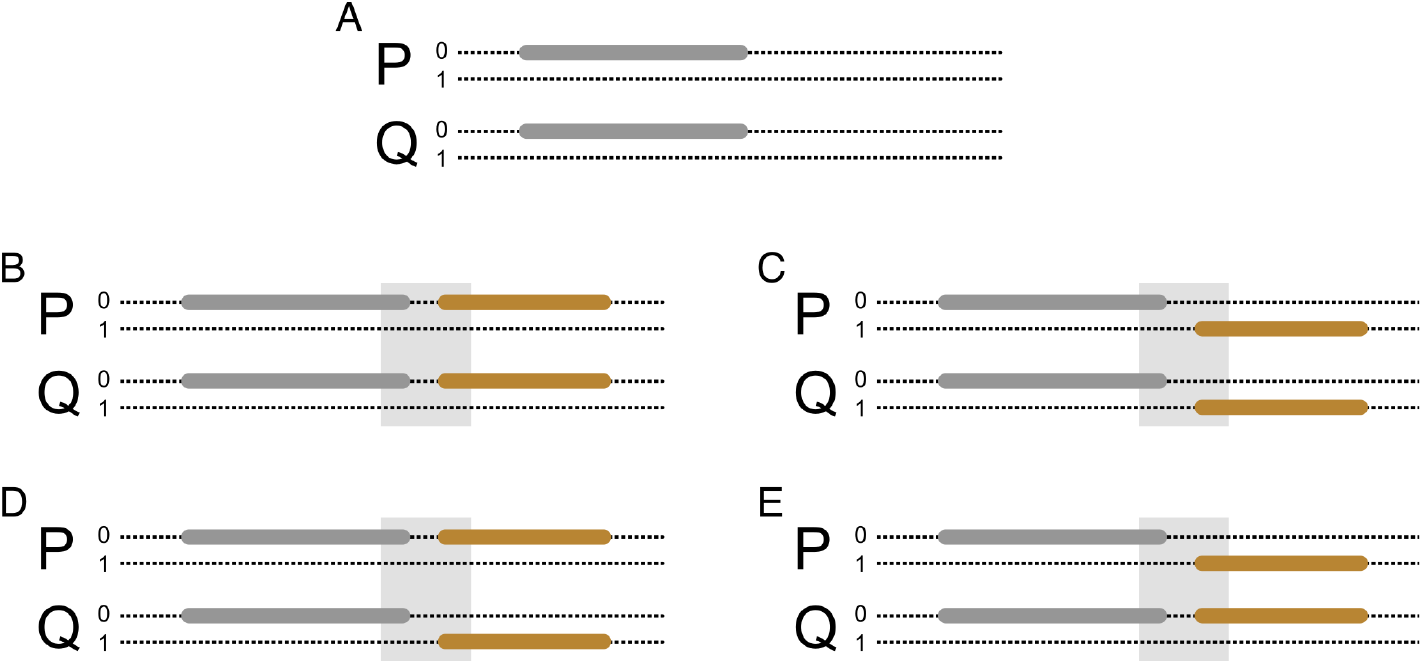
Possible scenarios considered by the TPBWT for adjacent IBD segments. IBD segments that slightly overlap or have a short gap between them may arise either through actual recombination patterns or phase switch errors. (*A*) Shown are the two haplotypes (0 and 1; dotted lines) of two related individuals (*P* and *Q*) for a single chromosome. An IBD segment shared by *P* and *Q* is shown in grey. (*B*) As the TPBWT scans left to right along the chromosome, another IBD segment (orange) is found. If the new segment begins within a small interval near the end of the old segment (light grey box) and the new segment is on the same haplotypes as the old segment, then possibly the two segments are fragments of a longer segment broken up by error and should be merged. (*C*) If the new segment begins near the end of the old segment and the new segment is *not* on the same haplotypes as the old segment, then possibly there was a phase switch error in both individuals. (*D*) If the new segment begins near the end of the old segment and the new segment is on the same haplotype as the old segment in individual *P* but the complementary haplotype in individual *Q*, then possibly there was a phase switch error in individual *Q*. (*E*) If the new segment begins near the end of the old segment and the new segment is on the same haplotype as the old segment in individual *Q* but the complementary haplotype in individual *P*, then possibly there was a phase switch error in individual *P*.

If a phase switch error has been identified in either individual *P, Q*, or both, then the TPBWT will swap the haplotypes for the individuals containing the error (Figure 11). Now the new IBD segments merge and extend the fragments on the complementary haplotype that were broken up by the phase switch error. When the arrangement of IBD segments on the complementary haplotypes again suggests another phase switch error has been encountered the algorithm stops swapping the individual’s haplotypes. This process continues to the end of haplotypes “stitching” short stretches of IBD fragmented by errors back into the correct long IBD segments. Note that in the TPBWT, when the complementary haplotypes of an individual are swapped due to a phase switch error the two haplotypes are swapped for all templates simultaneously.

In this way, information regarding errors identified using one template is shared with the other templates to improve phase correction and thus IBD detection overall. Additionally, as noted earlier, this means that phase switch errors are fixed consistently throughout the entire cohort; when a phase switch error is identified and corrected for an IBD segment shared between two individuals any other IBD segments shared with other individuals affected by the same switch error will also be corrected.

Gaps between subsegments are commonly caused by consecutive phase switch errors that fragment long IBD segments. If the distance between the consecutive phase switch errors is less than the length threshold needed to be considered an IBD subsegment, then the fragment of IBD will be dropped causing a gap. For this reason we merge subsegments that are separated by a distance less than that length (determined by parameter *L*_*m*_). Note that *L*_*m*_ is the minimum length threshold (in the number of sites) for a subsegment to be merged into the putative IBD segment stored in *P*_*s*_ and *P*_*e*_. The putative IBD segments in *P*_*s*_ and *P*_*e*_ must still exceed length *L*_*f*_ (in cM) if they are to be reported to the user as a “good” IBD segment.

### 4.3 TPBWT-Compressed Haplotypes

Durbin (2014) described how to leverage shared haplotype structure identified by PBWT to efficiently compress the haplotypes. At each position the haplotypes are sorted by the PBWT so that those with similar prefixes are adjacent to one another. Linkage disequilibrium causes correlation among sites close to one another on the chromosome and so haplotypes that share an allele at the current position will often share an allele at the next position. This creates long runs of the same allele in the PBWT sorted haplotype order which can be run-length encoded.

We use a similar run-length compression with the TPBWT. However, the compression scheme is slightly less efficient than with PBWT since at each position we may have multiple haplotype orderings that must be encoded. For example if site *k* is processed by three templates, than site *k* will have three haplotype orderings in *ppa*_*j,k*_ and so is run-length compressed three separate times. While this still results in significant file size reductions the primary benefit is that parsing a TPBWT-compressed haplotype file can be much faster than parsing other representations of the haplotypes. This is because in Algorithm 1 at site *k* the allele of each haplotype is queried when the haplotype is encountered in *ppa*_*j,k*_. If those alleles are already run-length encoded using the haplotype order *ppa*_*j,k*_ then we can modify Algorithm 1 so that alleles are only queried when they are at the beginning of a new allele run rather than for every haplotype. This can dramatically reduce the time needed to parse haplotypes from a file during an IBD analysis. However, since generating the TPBWT-compressed haplotypes takes the same time as computing the IBD shared among those haplotypes the TPBWT-compressed haplotypes are not necessarily useful for in-sample IBD analyses unless one is trying to save disk space. Rather we find that the TPBWT-compressed haplotypes are most useful for algorithms that utilize the TPBWT data structure to make estimates other than in-sample IBD, for example out-of-sample IBD analyses.

### 4.4 Out-of-Sample Analyses

A common application for the large sample size cohorts in biobank or DTC genetic databases is out-of-sample IBD computation, for example when comparing new samples to an existing large set of samples. For these analyses we use a modified form of Algorithm 1 in which the two haplotype alignments are essentially treated as one; the algorithm passes through both sets of samples at the same time and only reports the IBD segments shared among old and new samples. For this approach a major bottleneck is the memory used to store *𝒫*_*s*_ and *𝒫*_*e*_. If we have two sets of samples *X* and *Y*, each with *M*_*X*_ and *M*_*Y*_ haplotypes respectively, then *𝒫*_*s*_ and *𝒫*_*e*_ will be *M*_*X*_ + *M*_*Y*_ by *M*_*X*_ + *M*_*Y*_ in size. This would be prohibitive for very large sample sizes. However since we are only interested in the matches between *X* and *Y* and not the matches within either dataset we modify *𝒫*_*s*_ and *𝒫*_*e*_ to be *M*_*X*_ by *M*_*Y*_ in size, substantially reducing the memory required for out-of-sample analyses.

This algorithm can be highly efficient if both sets of samples have already been TPBWT-compressed. In this case, in our two sets of samples *X* and *Y* at site *k* we will already have run-length encoded the alleles according to two positional prefix arrays 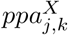 and 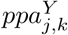, respectively. For the out-of-sample analysis we need to look up alleles ordered by the positional prefix array of the combined sample sets 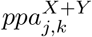. Here we take advantage of the fact that 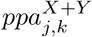 is the linear sum of the two totally ordered sets 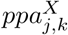 and 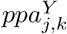. This means that within 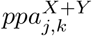 the haplotypes from *X* will follow the order found in 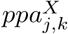 and the haplotypes from *Y* will follow the order found in 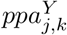. Instead of querying the allele for every single haplotype in 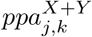 we now need to only query alleles if they are at the beginning of a new allele run encoded by 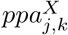 or 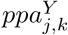.

### 4.5 Implementation

The TPBWT is available for non-commercial use as the Python package phasedibd in the code repository https://github.com/23andMe/phasedibd. It is implemented in Cython (Behnel et al. 2011) and compiles into both Python 2.7 and Python 3 (Van Rossum and Drake Jr 1995; Van Rossum and Drake 2009).

### 4.6 Parallelization For Large Sample Sizes

Our software implementation allows for a number of highly flexible parallelization schemes that enable fast and efficient IBD computes over extremely large cohorts. Scaling up to large sample sizes we use a simple batch method that utilizes TPBWT-compressed haplotypes. For each chromosome:

1. Divide the *M* haplotypes into *b* equally sized batches (one VCF file for each batch).
2. In parallel, on *b* CPU cores, compute the IBD shared among the haplotypes within each batch. During this compute write the TPBWT-compressed haplotypes to *b* binary files.
3. Use 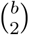 CPU cores to compute the out-of-sample IBD shared between batches. This utilizes the TPBWT-compressed haplotypes to increase the efficiency of each out-of-sample IBD compute.

See an example with compute times in the Results section.

Similar batching approaches are useful for running large out-of-sample analyses; for example when new samples have been acquired and must be run against a large panel of existing samples. If the existing samples have already been TPBWT-compressed in batches, the new samples can be easily compared to the existing samples in parallel. These massively parallel out-of-sample analyses over TPBWT-compressed haplotypes can result in substantial decreases in wall-clock compute time needed for biobank-scale data sets.

### 4.7 Simulation Study and Comparisons to Other Methods

To assess the accuracy of IBD inference methods we utilized both randomly sampled sets of genotyped research consented 23andMe customers and simulated haplotype data sets in which the IBD segments shared were perfectly known. For the simulated haplotypes we introduced realistic levels of genotyping and phasing errors to test the impact of these errors on inference.

#### 4.7.1 Simulating Haplotypes

We simulated haplotypes inherited with recombination over 400 replicated pedigrees. Each pedigree had three generations and included at least one pair of each type of close relatives that were used for the simulation study: parent-child, grandparent-grandchild, aunt-niece, first cousins, and siblings. Each pedigree founder consisted of a randomly sampled and unrelated research consented 23andMe customer. Recombination was simulated using a Poisson model with a rate of 1 expected crossover per 100 cM. This resulted in simulated haplotypes for 2000 closely related pairs of individuals with perfectly known IBD segments, 400 pairs of each relationship type: parent-child, grandparent-grandchild, aunt-niece, first cousins, and siblings.

#### 4.7.2 Simulating Genotyping Errors

We incorporated a simple model of genotyping error into our simulated data sets. At each position along the simulated chromosomes we introduced error into the genotype call with a probability of 0.001. When a site was selected for an error, half of the genotype call would be “flipped” with equal probability (e.g. a 0/0 genotype would be converted to a 1/0 or a 0/1 genotype with equal probability).

#### 4.7.3 Simulating Phasing Errors

We introduced errors due to statistical phasing into our simulated haplotype datasets. We first converted all the simulated haplotypes into their respective diploid genotypes and then used the statistical haplotype phasing method Eagle2 (Loh et al. 2016). For the phasing reference panel we used an internal 23andMe phasing panel that included about 200000 non-Europeans and about 300000 Europeans. This resulted in simulations that had a mean switch error rate of 0.25%, comparable to switch error rates measured elsewhere (Choi et al. 2018).

#### 4.7.4 Comparing Performance of TPBWT to Other Phase-Aware Algorithms

Table 2 outlines the parameter settings used for the different phase-aware methods. To avoid the possibility of erroneously conflating very short nearby IBD segments into long segments we only estimated IBD segments at least 3 cM or longer (Chiang et al. 2016) for all methods except Durbin’s PBWT which does not use genetic distance. PBWT requires the minimum number of sites in a segment to be specified; we set this to be 200 sites. TPBWT requires both a minimum segment length in genetic distance and a minimum number of sites; we set these to be 3.0 cM and 200 sites, respectively. The same parameter settings were used in all comparative analyses.

To compare the accuracy of IBD estimates made by each phase-aware method we used the simulated datasets described above and calculated the error in two summary statistics: the proportion of the genome that is IBD between two individuals and the number of IBD segments shared among the two individuals. These two statistics are particularly informative when estimating relatedness or other demographic quantities from IBD segments. We calculated the percent of the genome that was erroneously inferred to be IBD for a simulated pair of close relatives as 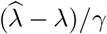 where *λ* is the true total amount of the genome that is IBD, 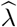 is the estimated amount of the genome that is IBD, and *γ* is the genome length. We calculated the number of erroneous IBD segments estimated for a simulated pair of closes relatives as 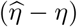 where *η* is the true number of IBD segments and 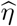 is the estimated number of IBD segments.

To further compare the methods’ performance we additionally calculated false positive and false negative rates of inferring IBD segments by their length. Rates were calculated for bins of IBD segment lengths: 3–4, 4–5, 5–6, 7–8, 9–10, 10–11, 12–15, 15–18, and *>* 18 cM. To thoroughly explore these rates and their effects on IBD estimates, we calculated each rate in two different ways. *False negative rate by segment* is the proportion of true segments in a size bin that do not overlap any estimated segment compared to the total number of true segments in the size bin. *False negative rate by segment coverage* is the proportion of the length of true segments in a size bin not covered by any estimated segment compared to the total length of true segments in the size bin. *False positive rate by segment* is the proportion of estimated segments in a size bin that do not overlap any true segment compared to the total number of true segments in the size bin. *False positive rate by segment coverage* is the proportion of the length of estimated segments in a size bin not covered by any true segment compared to the total length of true segments in the size bin.

To compare the computation time needed for each phase aware method we randomly sampled sets of research consented 23andMe customers genotyped on the 23andMe v5 microarray chip. We removed SNPs with *<* 85% genotyping rate, SNPs with MAF *<* 0.001, SNPs with low trio concordance (effect *<* 0.6 and p-value *<* 1e-20), and SNPs with allele counts of 0 within the samples selected for the phasing reference panel. After this quality control filtering a total of 544,042 SNPs were used. Haplotypes were phased using Eagle2 (Loh et al. 2016) with a reference panel containing 286,305 samples. IBD was computed for 42,927 SNPs from human chromosome 1. Though iLASH, hap-IBD, and Refined IBD are multithreaded programs the runtimes compared in this test were for single-threaded operation on a single CPU core. Note all IBD inference can be trivially parallelized using batching approaches (see Section 2.3 and Table 1).

#### 4.7.5 Comparing Performance of TPBWT to a Phase-Free Algorithm

We compared the performance of TPBWT to the IBIS-like IBD inference algorithm that uses unphased data. To compare the accuracy of detecting IBD with the IBIS-like algorithm and the TPBWT, and since unphased approaches are expected to have higher false positives especially on shorter segments, we replicated the trio validation approach used in Henn et al. (2012). We randomly sampled 1,000 child-parent trios and 10,000 individuals not in any of the trios from research consented 23andMe customers. Each customer was genotyped on the 23andMe v5 microarray chip. We removed SNPs with *<* 85% genotyping rate, SNPs with MAF *<* 0.001, SNPs with low trio concordance (effect *<* 0.6 and p-value *<* 1e-20), and SNPs with allele counts of 0 within the samples selected for the phasing reference panel. After this quality control filtering a total of 544,042 SNPs were used. We computed IBD among all 13,000 individuals using TPBWT and IBIS-like. For the TPBWT compute the haplotypes were phased using Eagle2 (Loh et al. 2016) with a reference panel containing 286,305 samples. Since both Henn et al. (2012) and Seidman et al. (2020) showed that IBIS-like algorithms have high false positive rates for segments *<* 7 cM in length, we used 7 cM has the minimum segment length for the IBIS-like algorithm.

For each observed IBD segment shared between a child and a distant relative, we labeled each segment as either “trio validated” or “not trio validated”. Segments were trio validated if an overlapping segment was observed to be shared between the distant relative and one or more of the child’s parents. Segments were not trio validated if no overlapping segment was found between the child’s parents and the distant relative. For bins of IBD segment lengths we then calculated *h*_mean_, which is the proportion of IBD segments in that length bin that were trio validated. Segments that were not trio validated were either false positive segments in the child or false negative segments in the parents.

### 4.8 Case Study: Haplotype Sharing in Mexico

To demonstrate the utility of the IBD estimates made using the TPBWT and the 23andMe database we performed a brief case study to examine the geographic patterns of haplotype sharing within Mexico. We identified 9,517 research consented 23andMe customers who self reported that all 4 of their grandparents were from the same Mexican state. Each customer was genotyped on either the 23andMe v4 or v5 microarray chip. We removed SNPs with *<* 85% genotyping rate, SNPs with MAF *<* 0.001, SNPs with low trio concordance (effect *<* 0.6 and p-value *<* 1e-20), and SNPs with allele counts of 0 within the samples selected for the phasing reference panel. After this quality control filtering the v4 chip had 453,065 SNPs and v5 chip had 544,042 SNPs. Haplotypes were phased using Eagle2 (Loh et al. 2016). Individuals on the v4 chip were phased with a reference panel containing 691,759 samples. Individuals on the v5 chip were phased with a reference panel containing 286,305 samples.

IBD sharing among the 9,517 individuals was computed using the TPBWT with the parameters described in Table 2. IBD estimates among individuals on the same genotyping chip were made using the in-sample method described above, and estimates made among individuals on different chips was made using the out-of-sample approach described above over the intersection of chip SNPs (only the SNPs present in both the v4 and v5 genotyping chips). Hierarchical clustering of the mean pairwise IBD haplotype sharing across Mexican states was performed using Ward’s method (Ward Jr 1963) in R (R Core Team 2013). To remove close relatives we excluded any pair of individuals that shared more than 20 cM. Geographic maps of the mean pairwise IBD shared across Mexican states were made using the R packages mxmaps, ggplot2, and viridis (Valle-Jones 2019; Wickham 2016; Garnier 2018).

## 5 Data Availability

The data underlying this article cannot be shared publicly to protect participant privacy and in accordance with the IRB-approved protocol under which the study was conducted. Upon request, de-identified summary statistics will be shared for use in research through a data transfer agreement.

## 6 Acknowledgements

We thank Richard Durbin for insightful feedback and inspiration. Also we thank Dave Hinds, Steven Micheletti, David Poznik, and Sayantan Das for providing useful comments. Additionally, we thank two anonymous reviewers for detailed comments that significantly improved the manuscript. This work would not have been possible without the 23andMe customers who consented to participate in research – thank you. And finally we thank all the employees of 23andMe who contributed to the development of the infrastructure that made this research possible.

